# Distinct ‘pattern of autofluorescence’ of acute ischemic stroke patients’ skin and fingernails: A novel diagnostic biomarker for acute ischemic stroke

**DOI:** 10.1101/310904

**Authors:** Danhong Wu, Mingchao Zhang, Yue Tao, Haibo Yu, Yujia Li, Shufan Zhang, Tianqing Chu, Xiaofeng Zou, Xin Chen, Weihai Ying

**Author notes:** Corresponding author: Weihai Ying, Ph.D. Professor, School of Biomedical Engineering and Med-X Research Institute Shanghai Jiao Tong University 1954 Huashan Road Shanghai, 200030, P.R. China. These authors contributed equally to this work (Danhong Wu, Mingchao Zhang, Yue Tao).

## Abstract

Early and economical diagnosis of acute ischemic stroke (AIS) is pivotal for therapeutic efficacy, particularly for the settings where medical imaging resource is deficient. We have obtained evidence supporting our hypothesis that collective properties of the green autofluorescence (AF) of the fingernails and certain skin’s positions may be a novel diagnostic biomarker for AIS: Both the green AF intensity and AF asymmetry of the AIS patients in their Index Fingernails and most examined skin’s positions were significantly higher than that of the healthy subjects and the Non-AIS subjects. ROC analyses and machine learning-based analyses on the AF properties showed that AUC was 0.93 and 0.87, respectively, for differentiating the AIS patients from the healthy subjects and for differentiating the AIS patients from the Non-AIS subjects. The AIS patients had significantly higher AF intensity and AF asymmetry at several examined positions, compared to those of the patients of Parkinson’s disease, pulmonary infection and transient ischemic attack. The AUC was 0.79 – 0.88 for differentiating AIS patients from each of these diseases. There was evidence suggesting that the AF originates from keratins. Collectively, our study has indicated that the characteristic AIS’s ‘Pattern of AF’ is a novel diagnostic biomarker for the disease. The ‘Pattern of AF Technology’ holds excellent potential to become a new non-invasive, label-free and economical diagnostic approach for AIS, which is particularly valuable when MRI or CT imaging resource is deficient.

## Introduction

Stroke is one of the leading causes of death and disability around the world^1^. The only FDA-approved drug for treating ischemic stroke – recombined tissue plasminogen activator (rt-PA) - has very limited use, mainly due to its short therapeutic window ^2^: The drug must be administered within 4.5 hours after the incidence of acute ischemic stroke (AIS)^3^, otherwise therapeutic effects would be diminished and major detrimental effects such as hemorrhagic transformation may ensue. AIS patients with large artery occlusion can also be treated by Endovascular therapy with mechanical thrombectomy within 6 hours of symptom onset^4^. However, from the time of incidence of AIS, it usually takes greater than 6 hours for an AIS patient to get diagnosis. It was calculated that an acute stroke patient losses 1.8 million neurons every minute without appropriate treatment^5^. Therefore, early diagnosis has become a key factor that determines therapeutic effectiveness for the disease. To solve this critical problem, it has become increasingly important to establish novel approaches that can conduct AIS diagnosis non-invasively and efficiently before patients enter clinical settings.

Another important problem that limits efficient diagnosis of AIS patients has originated from the deficiency of medical imaging resource: MRI and CT imaging plays crucial roles in AIS diagnosis^6, 7^. However, there has been significant deficiency in MRI and CT imaging resource to meet the urgent demand of AIS diagnosis in many places around the globe, particularly in developing countries^8, 9^. Even in developed countries, MRI imaging resource is deficient in acute settings^10^.

The AF of the advanced glycation end-products (AGEs) of collagen and elastin in the dermis has shown promise for non-invasive diagnosis of diabetes and diabetes-associated diseases ^11–13^. Our latest study has reported that ROC analyses using the skin’s green AF intensity at Dorsal Centremetacarpus as the variable showed that the area under curve (AUC) for differentiating lung cancer patients from pulmonary infection (PI) patients was 0.87^14^. Since AF detection is non-invasive, economical and rapid, it is of great significance to further investigate the potential of AF as diagnostic biomarkers for diseases.

In our current study, we tested our hypothesis that AIS patients may have characteristic green AF properties in their fingernails and certain skin’s positions, which may become a novel diagnostic biomarker for AIS. Our study has provided multiple lines of evidence supporting this hypothesis.

## Methods and materials

### Human subjects

The studies on the group of healthy subjects (Healthy Group), the group of AIS patients (AIS Group), the group of Non-AIS patients (Non-AIS Group), the group of the persons with high risk for developing AIS (High-Risk Group), the group of ischemic stroke patients in recovery phase (Recovery Group), the group of Parkinson’s disease group (PD Group), the group of PI (PI Group), and the group of transient ischemic attack (TIA Group) were conducted according to protocols approved by the Ethics Committee of Shanghai Fifth People’s Hospital Affiliated to Fudan University and the Ethics Committee of Shanghai Chest Hospital Affiliated to Shanghai Jiao Tong University. The following statement is the inclusion and exclusion criteria for the human subjects in each group: The subjects of the Healthy Group were selected through the screening of the community populations of the City of Shanghai based on the following criteria: (1) Aged 50 - 80; (2) with zero risk factors; (3) without previous AIS or current cerebral hemorrhage; (4) without neoplastic diseases and severe liver or kidney dysfunctions; (5) without autoimmune diseases or infection at active phase; and (6) without hypothyroidism. The ischemic stroke study was conducted from January 1, 2018 to April 30, 2019. Patients who met the following criteria were included in the AIS group: (1) MRI-proven AIS; (2) symptom onset within 7 days; and (3) aged 50 - 80. Patients were excluded if they met the following criteria: (1) Patients with previous AIS or current cerebral hemorrhage; (2) neoplastic diseases, and severe liver or kidney dysfunctions; (3) autoimmune diseases or infection at active phase; and (4) hypothyroidism. The subjects of the Non-AIS Group had the same exclusion criteria as those for the AIS patients. The patients who met the following criteria were included in the Non-AIS group: (1) Patients who were hospitalized in the Department of Neurology, Shanghai Fifth People’s Hospital Affiliated to Fudan University from January 1, 2018 to April 30, 2019; and (2) the diagnostic results from both CT and DWI imaging were negative. The subjects of the High-Risk Group had the same exclusion criteria as those for the AIS patients. The subjects of the High-Risk Group met the following criteria: (1) The diagnostic results from both CT and DWI imaging were negative; (2) the subjects had at least three risk factors out of the following seven risk factors: Hypertension, diabetes, hyperlipidemia, atrial fibrillation, body mass index (BMI) > 26, family history of stroke, and smoking; and (3) the subjects were not TIA patients. The inclusion and exclusion criteria for the subjects of the Recovery Group were the same as those of the AIS Group, except that the subjects of the Recovery Group had AIS incidence 1 - 3 months before the AF determinations. The subjects of the PD Group met the following criteria: (1) aged 50 – 80, (2) met the UK idiopathic Parkinson’s disease brain bank criteria^15^, and (3) measured by 24-h ABPM. Patient exclusion criteria was the same as those for the AIS group. The study on the PI group was conducted from January 1, 2018 to June 30, 2018. All patients were aged between 50-80 and met the criteria of the ‘Guidelines for the diagnosis and treatment of community-acquired pneumonia in adults in China’ ^16^. The baseline information of each group of the subjects was shown in Supplemental Table 1.

### Development of A Portable AF Imaging Device and Image Processing Program

The portable AF imaging device consists of an imaging system and a LED excitation apparatus. The imaging system consists of the lens (U-TLU, Olympus) and optical filters (ET4845/10, Chroma; ET525/50, Chroma) which are coupled with a CMOS camera (acA1920-50gm, Basler). When the imaging system is operating, a LED (PL35-8-5, Couson) is employed as the excitation source for the AF imaging at the excitation wavelength of 485 nm ± 10 nm. The light with emission wavelength of 500 – 550 nm which is filtered through the optical filter is received by the CMOS camera. The images from the CMOS camera are processed through an imaging software at an external computer terminal. The parameters of the devices and the protocols for the AF imaging are stated in supplementary information. By using a portable device that was made by combining a spectrometer (USB4000, Ocean Optics) and the optical apparatus of the portable AF imaging device, the spectra of the skin’s AF of human subjects were determined. The emitting light of the wavelength longer than 450 nm was detected, when the excitation wavelength of 445 nm was used.

### The equipment parameters and imaging protocols

The following parameters of the device were used for determinations of the green AF of the skin and fingernails of the human subjects: Laser power in working state: 9.79 μW; microscopy type: widefield; exposure time: 22000 μs; frame rate: 100 frames/second; imaging resolution:1450 pixels/mm; image size: 1920 * 1200 pixels; physical size: 1.32 mm * 0.83mm.

To prevent the interferences of experimental conditions and other variables on the experimental results, the following protocols were conducted: 1. The protocols for preventing the interferences of environmental factors on the experimental results: (1) The measurements should be conducted in dark room to avoid the interference of background light on the AF readings; (2) room temperature is maintained at 20 - 24℃, and the humidity is 40% - 60%; and (3) the subjects are not disturbed by environmental factors. 2. The protocols for preventing the interferences of the hygiene conditions of the subject’s skin and fingernails on the experimental results: (1) The subject’s skin and fingernails should be cleaned by alcohol-free water. Measurements should not start until 10 – 20 min after the water on the skin and fingernails is removed; (2) no chemicals can be applied to the skin or fingernails 12 hours before the measurements; and (3) the subject’s skin and fingernails is cleaned by alcohol swabs carefully before the measurements. 3. The protocols for preventing the interferences of the healthy conditions of the subjects on the experimental results: The subjects with the following medical conditions should be excluded from the study: (1) the subjects with pathological conditions of their skin; (2) the subjects who have been treated for their skin diseases within one month; (3) the subjects who have diseases that may affect their skin conditions; (4) the subjects who have undergone immunosuppression therapy within 3 months; (5) the subjects who have undergone systemic hormone therapy or phototherapy within one month; and (6) the subjects who have undergone therapies with applications of anti-itch medications, sex hormones, anti-depression drugs, immunomodulation drugs, or exfoliating agents. 4. To assure the stability of the equipment for measuring the AF of the skin and the fingernails of subjects, the laser power is measured regularly to assure the laser power is stable throughout the period of the study.

### Determinations of AUC by machine learning-based analyses on the AF images

The AF images of the left and right Ventriantebrachium, Dorsal Antebrachium, Dorsal Centremetacarpus, Centremetacarpus, Dorsal Index Fingers, Ventral Forefinger and Index Fingernails were utilized to differentiate the AIS patients from the healthy subjects, the Non-AIS subjects, the PD patients, the PI patients and the TIA patients.

The following was the image feature extraction method: Took a square window with a side length of 25, scanned the image with a step length of 5, and calculated the average value of the brightness of all points in the window as the value of the window. The maximum value was taken as the eigenvalue. In order to fully extract the global information of the image, the window completely falling in the square area with the maximum window as the center and the side length of 275 was removed. Repeated these operations for ten times, and ten initial eigenvalues were extracted from each image. The datasets were split into training sets and testing sets by stratified k-fold method (k=3). For each set of data, the first phase was feature selection based on the training set. To verify the robustness, another k-fold cross-validation was performed, and LightGBM algorithm was adopted. Calculated the average value of k-time feature importance and classification accuracy. The features were retained from large to small according to the feature importance. When the sum of the feature importance of the retained features was greater than or equal to 95% of the sum of the feature importance of all features, the model was trained with the retained features. If the classification accuracy was improved, these operations were repeated. If the prediction result became worse, all the features of the previous round were considered as input features. Subsequently, hyperparameter combination selection of the model was conducted, which was chosen according to different combination’s performance of stratified k-fold cross-validation on the train set. The model employed was also LightGBM classifier. The model was then trained on the training set to obtain model parameters. The model’s performance on the testing set was then evaluated with AUC, Recall Score, Precision Score, and F1 Score as index, using the same input features and hyperparameter combination as the training set. This process was repeated k times. Final outputs were determined by calculating the average values and standard deviations of the evaluation results. The machine learning pipeline is shown as the following diagram.

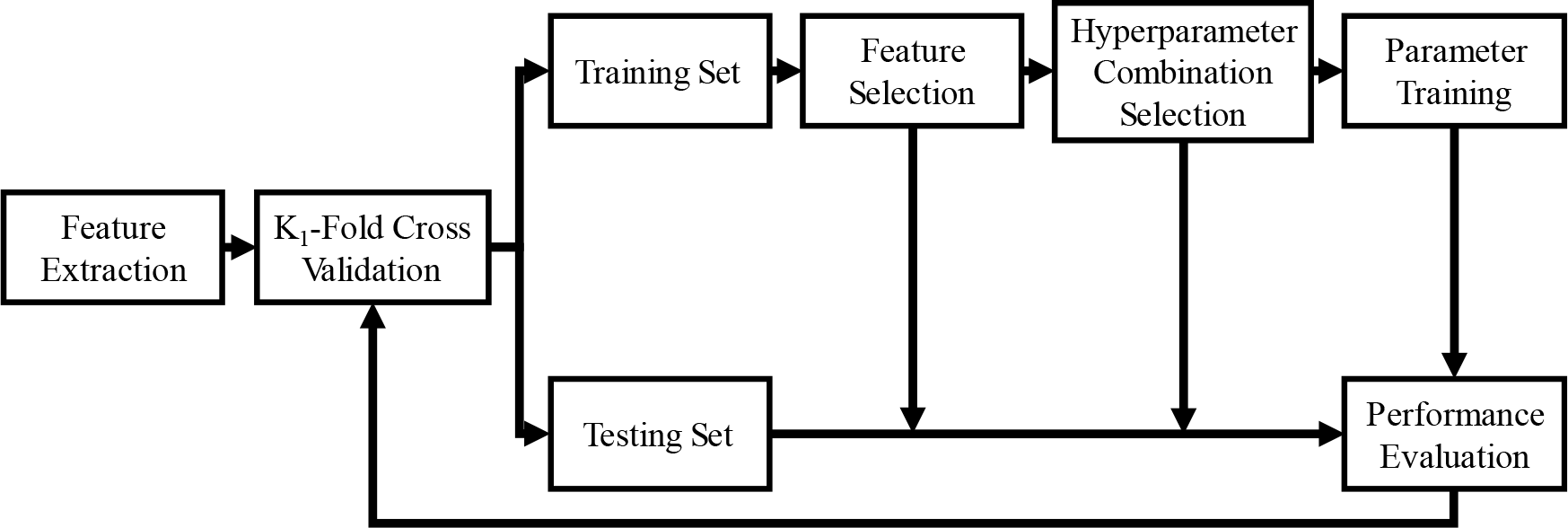

### Statistical analyses

All data were presented as mean ± SEM, except where noted. The data from the studies on human subjects were assessed by Kruskal-Wallis multiple comparison test (K-W test) or Mann-Whitney test. Graphpad prism 7.0 was used for both ROC analyses and determinations on if the data were normally distributed. *P* values less than 0.05 were considered statistically significant.

## Results

### 1. AIS patients had significantly higher green AF intensity at their Index Fingernails and most examined skin’s positions than that of the healthy subjects and the Non-AIS subjects

We determined the AF spectra of all examined positions of the healthy subjects, the Non-AIS subjects and the AIS patients: The AF spectra of the Index Fingernails and each examined skin’s position of the AIS patients were highly similar with each other, which matched that of keratins or FAD (Fig. 1A)^17, 18^. The AF spectra of the Index Fingernails and each examined skin’s position of the healthy subjects and those of the Non-AIS patients were also highly similar with each other, which matched that of keratins or FAD (Fig. 1A)^17, 18^. Based on the properties of the AF spectra, we determined the AF of the AIS patients, the healthy subjects and the Non-AIS patients by using the excitation wavelength of 485±10 nm, and the emission wavelength of 500 – 550 nm: At the Index Fingernails (Fig. 1B) and/or certain examined skin’s positions, AIS patients had higher AF intensity in a great majority of time, compared to that of the healthy subjects or the Non-AIS patients.

**Fig. 1.**
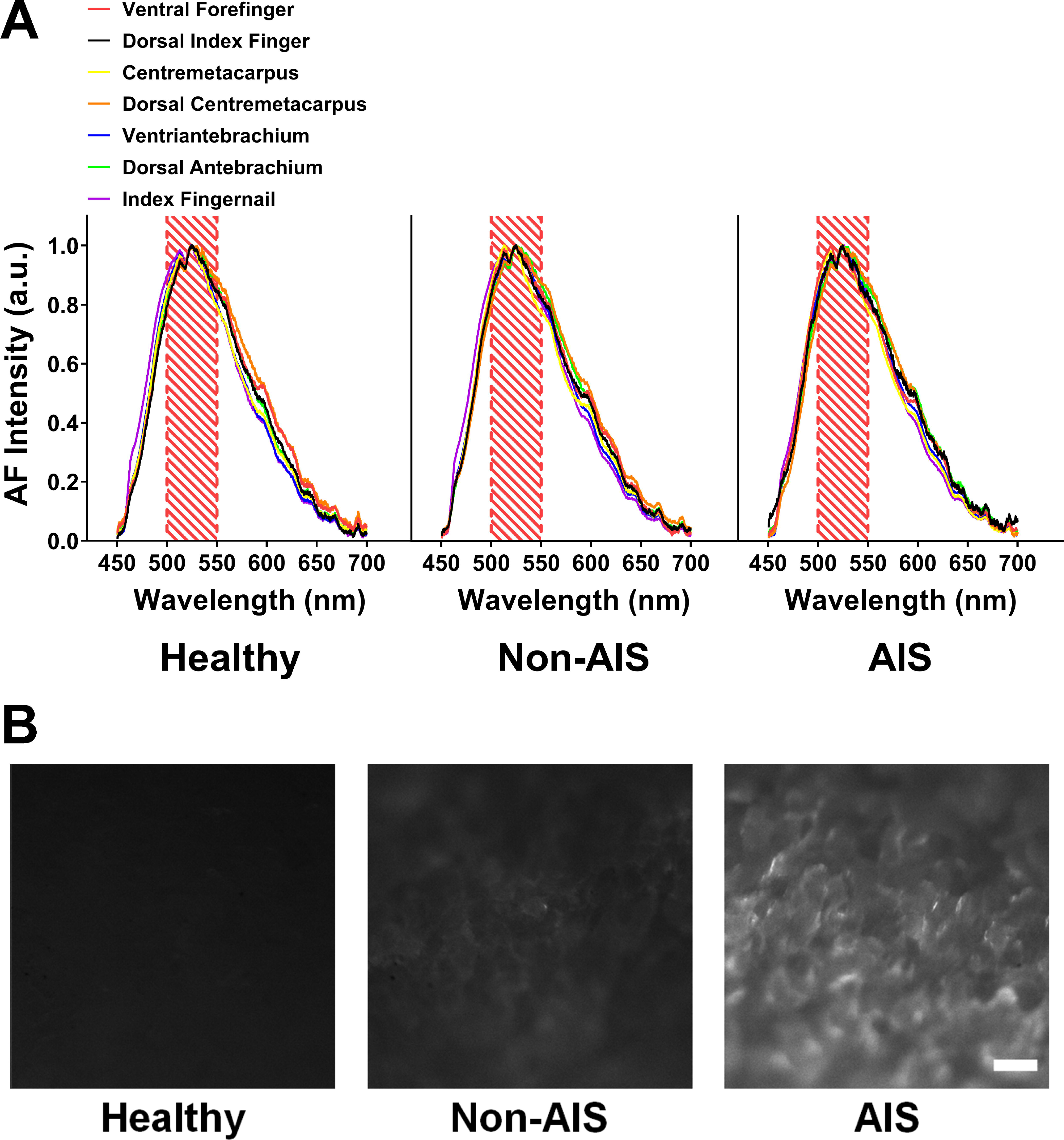
The AF spectra of each examined skin’s position and the Index Fingernails of the AIS patients matched that of keratins or FAD. (A) The AF spectra of the Index Fingernails and each of the twelve examined skin’s positions of the AIS patients were highly similar with each other, which matched that of keratins or FAD. The AF spectra of the Index Fingernails and each of the twelve examined skin’s positions of the healthy subjects and the Non-AIS subjects were also highly similar with each other, which matched that of keratins or FAD. The number of the healthy subjects, the Non-AIS subjects and the AIS subjects was 3, 3, and 3. (B) At the Index Fingernails, AIS patients had higher AF intensity in a majority of time, compared to that of the healthy subjects and the Non-AIS patients. The figures were representatives of the AF images of the Index Fingernails of the healthy subjects, the Non-AIS subjects and the AIS subjects. The number of the healthy subjects, the Non-AIS subjects and the AIS subjects was in 55, 117-122, 78-80, respectively.

Since the AF intensity at the right and left side of the examined positions of a majority of the AIS patients was distinctly different, we defined the difference (in absolute value) between the AF intensity of the right and left side of a subject as ‘AF Asymmetry’, with the side of the skin with higher AF intensity being defined as ‘Stronger AF Side’. We determined the AF intensity of the right side, the left side and the ‘Stronger Side’ of the examined skin’s positions and the Index Fingernails of the healthy subjects, the Non-AIS subjects and the AIS patients: At the ‘Stronger Side’, the right side and the left side of most examined skin’s positions and the Index Fingernails, the AF intensity of the AIS patients was significantly higher than that of the healthy subjects and the Non-AIS subjects, except at the left Dorsal Centremetacarpus (Fig. 2, A to G). In particular, the AF intensity of the ‘Stronger Side’ of the Index Fingernails and the Dorsal Index Fingers of the AIS patients was 341 – 386 % of that of the healthy subjects (Fig. 2, B and G).

**Fig. 2.**
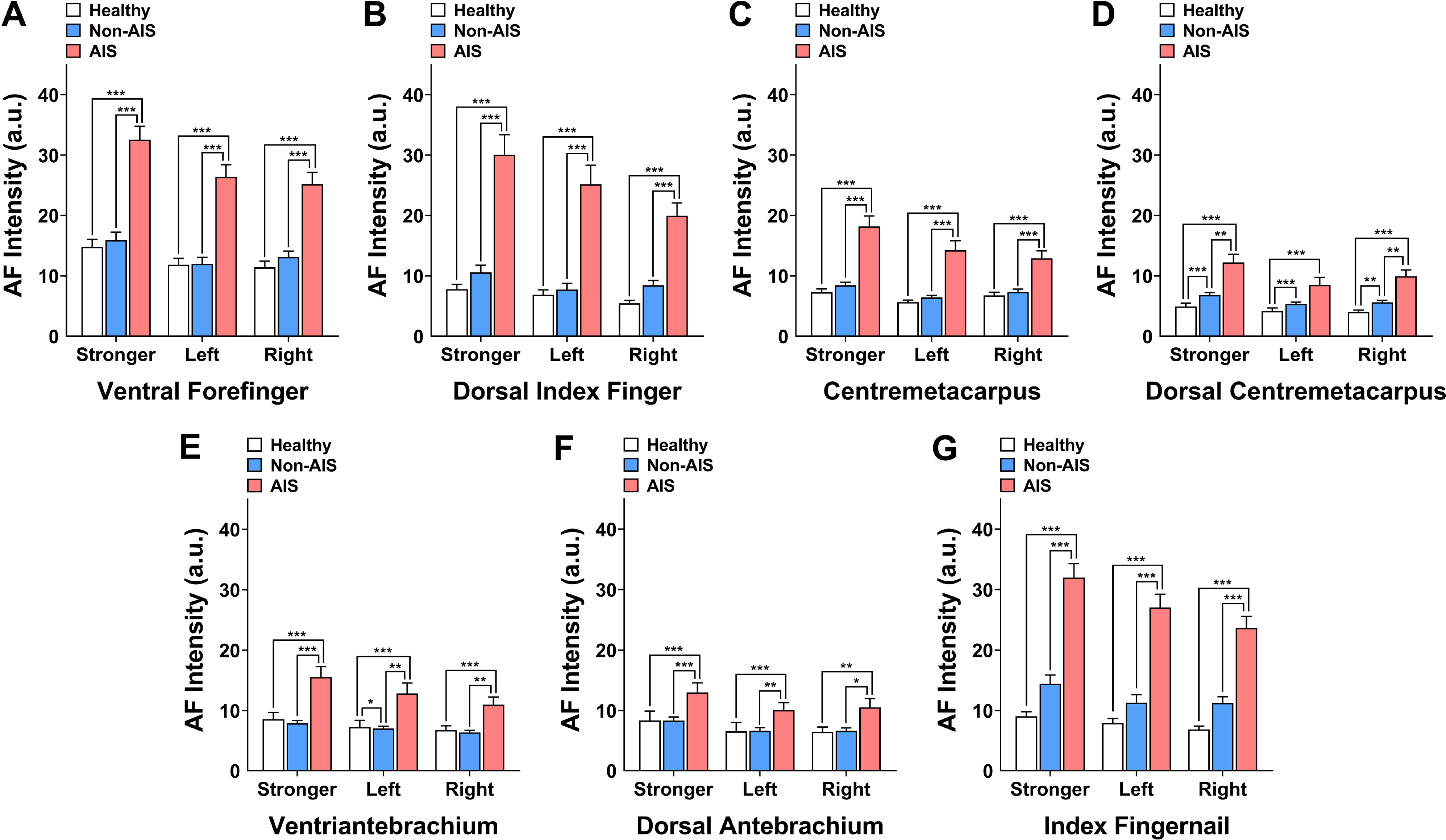
AIS patients had significantly higher green AF intensity at the Index Fingernails and most examined skin’s positions than that of the healthy subjects and the Non-AIS subjects. (A-G) At the Stronger Side, the right side and the left side of all examined skin’s positions and the Index Fingernails, the AF intensity of the AIS patients was significantly higher than that of the healthy subjects and the Non-AIS subjects, exception at the left Dorsal Centremetacarpus. The number of the healthy subjects, the Non-AIS subjects and the AIS subjects was in 55, 117-122, 78-80, respectively. *, *p* < 0.05; **, *p* < 0.01; ***, *p* < 0.001.

We also determined the green AF intensity of the High-Risk subjects at all examined positions: The AF intensity of the High-Risk subjects was significantly higher than that of the healthy subjects only at right Index Fingernails, the skin of the Stronger Side and the right side of Dorsal Index Fingers, and the Stronger Side, the left side, and the right side of Dorsal Centremetacarpus (Fig. S1, A to G). The AF intensity of the High-Risk subjects was significantly lower than that of the AIS patients at all examined positions, except at the skin of the Stronger Side, the left side and the right side of Dorsal Centremetacarpus, the skin of the left and the right Antebrachium, and the skin of the right Ventriantebrachium (Fig. S1, A to G).

The subjects of the Recovery Group had significantly lower AF intensity at all examined skin’s positions and the Index Fingernails than that of AIS patients, except at the skin’s positions of the left and right Centremetacarpus, right Ventriantebrachoum, right Dorsal Antebrachium, and the Stronger Side and the left side of Dorsal Centremetacarpus (Fig. S2, A to G).

### 2. AIS patients had significantly higher AF asymmetry at the Index Fingernails and all examined skin’s positions than that of the healthy subjects and the Non-AIS subjects

At all examined skin’s positions and the Index Fingernails, the AIS patients had significantly higher AF asymmetry than that of the healthy subjects and the Non-AIS subjects (Fig. 3, A to G). At the Index Fingernails and the skin’s positions of Dorsal Index Fingers, Dorsal Centremetacarpus and Centremetacarpus, the AF asymmetry of the AIS patients was 400 – 464 % of that of the healthy subjects (Fig. 3, A, C, D and G).

**Fig. 3.**
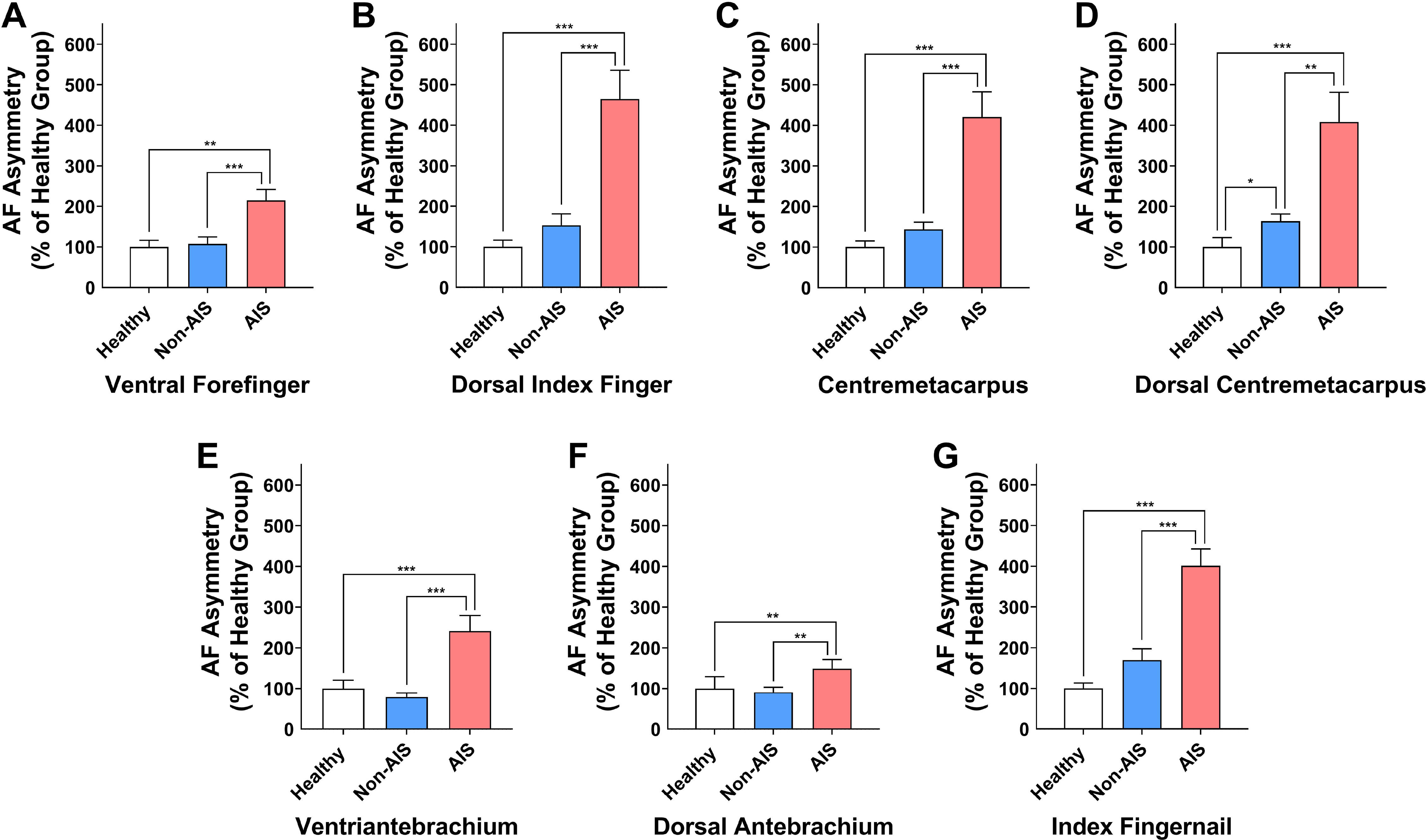
AIS patients had significantly higher AF asymmetry at the Index Fingernails and all examined skin’s positions than that of the healthy subjects and the Non-AIS subjects. (A-G) At all examined skin’s positions and the Index Fingernails, the AIS patients had significantly higher AF asymmetry than that of the healthy subjects and the Non-AIS subjects. The number of the healthy subjects, the Non-AIS subjects and the AIS subjects was in 55, 116-122, 67-80, respectively. *, *p* < 0.05; **, *p* < 0.01; ***, *p* < 0.001.

We also determined the AF asymmetry of the High-Risk subjects: The AF asymmetry of the High-Risk subjects was not significantly different from that of the healthy subjects at all examined positions, except at the skin’s position of Dorsal Centremetacarpus (Fig. S3, A to G). The AF asymmetry of the High-Risk subjects was significantly lower than that of the AIS patients at all examined positions, except at the skin’s positions of Centremetacarpus and Dorsal Antebrachium (Fig. S3, A to G).

### 3. AIS patients had markedly higher percentages of the subjects with four or more examined positions with increased AF intensity compared to those of the healthy subjects, the Non-AIS subjects and the High-Risk subjects

We determined the probability of each examined position with increased AF intensity for the healthy subjects, the Non-AIS subjects and the AIS subjects: Higher percentages of the AIS patients had increased AF at most examined positions, compared to those of the healthy subjects and the Non-AIS subjects (Fig. 4A). Particularly at right and left Index Fingernails, Dorsal Index Fingers and Ventral Forefingers as well as right Dorsal Centremetacarpus, the AIS patients had markedly higher probability to have increased AF intensity, compared to that of the healthy subjects and the Non-AIS subjects (Fig. 4A).

**Fig. 4.**
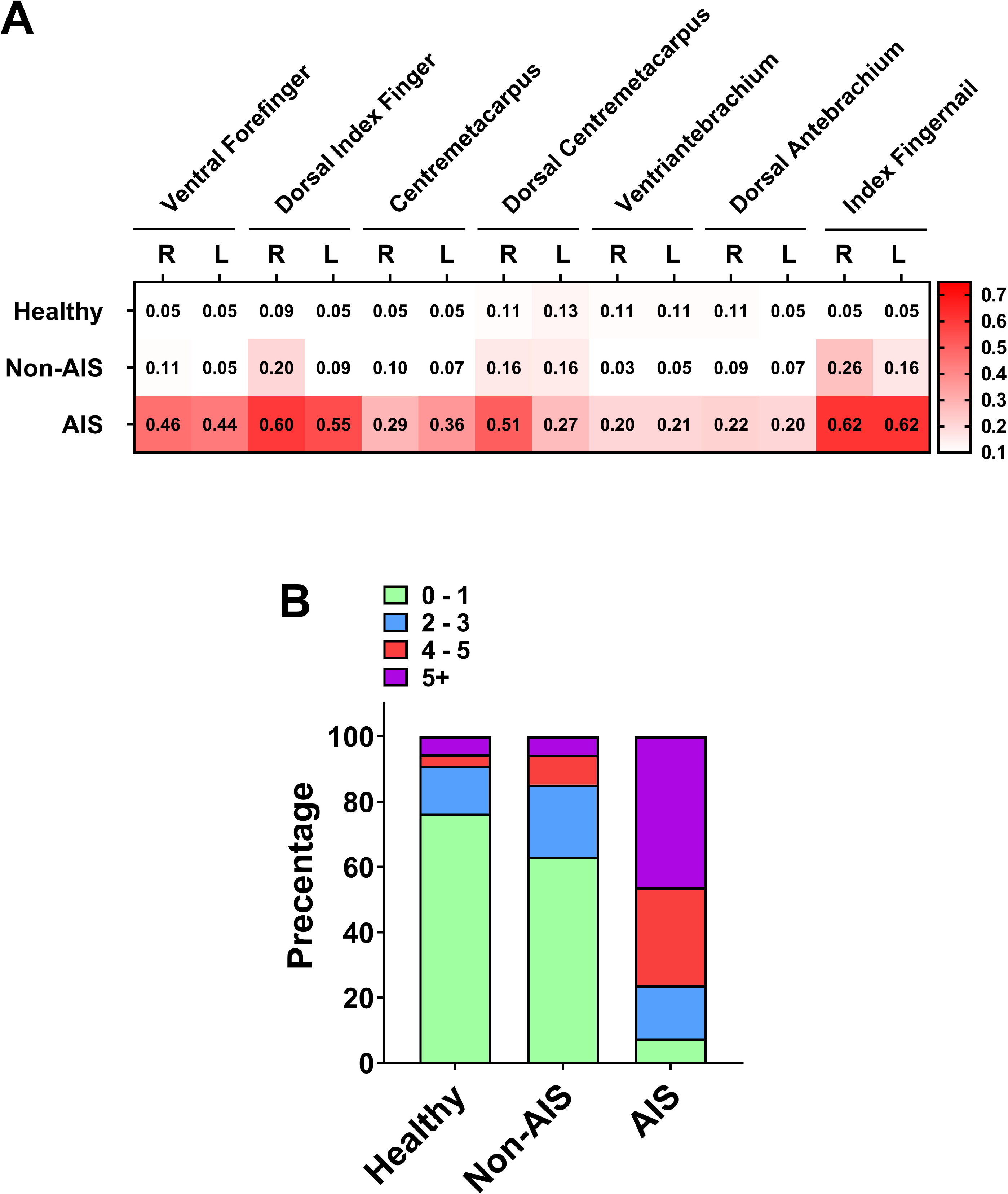
AIS patients had markedly higher percentages of the subjects with four or more examined positions with increased AF intensity compared to those of the healthy subjects and the Non-AIS subjects. (A) Higher percentages of the AIS patients had increased AF at most examined positions, compared to those of the healthy subjects and the Non-AIS subjects. (B) The percentage of the AIS patients with 4-5 or > 5 positions with increased AF intensity of the AIS group was 30% and 46%, respectively. In contrast, the percentage of the healthy subjects with 4-5 or > 5 positions with increased AF intensity was 4% and 5%, respectively; and the percentages of the Non-AIS subjects with 4-5 or > 5 positions with increased AF intensity of was 9% and 6%, respectively. The number of the healthy subjects, the Non-AIS subjects and the AIS subjects was 55, 122, 80, respectively.

We also determined if the AIS patients had markedly higher percentages of the people with four or more examined positions with increased AF intensity compared to those of the healthy subjects and the Non-AIS subjects: The percentage of the AIS patients with 4-5 or > 5 positions with increased AF intensity was 30% and 46%, respectively (Fig. 4B). In contrast, the percentage of the healthy subjects with 4-5 or > 5 positions with increased AF intensity was 4% and 5%, respectively (Fig. 4B); and the percentage of the Non-AIS subjects with 4-5 or > 5 positions with increased AF intensity of was 9% and 6%, respectively (Fig. 4B).

Higher percentages of the High-Risk subjects had increased AF at certain examined positions, compared to those of the healthy subjects (Fig. S4A). The percentage of the High-Risk subjects with 4-5 or > 5 positions with increased AF intensity was 14% and 14%, respectively (Fig. S4B).

### 4. ROC analyses and machine learning-based analyses on the AF properties showed significant promise of the AF-based diagnostic method for AIS

By using the green AF intensity at the Stronger Side, the left side or the right side of each examined position or using the AF asymmetry as the variable, ROC analyses were conducted to determine the AUCs for differentiating the AIS patients from the healthy subjects (Fig. 5, A and B). In particular, by using the AF intensity of the Stronger Side of the Dorsal Index Fingers, the Index Fingernails or the Ventral Forefingers as the variable, the AUC for differentiating the AIS patients from the healthy subjects was 0.88, 0.88 and 0.83, respectively (Fig. 5, A and B).

**Fig. 5.**
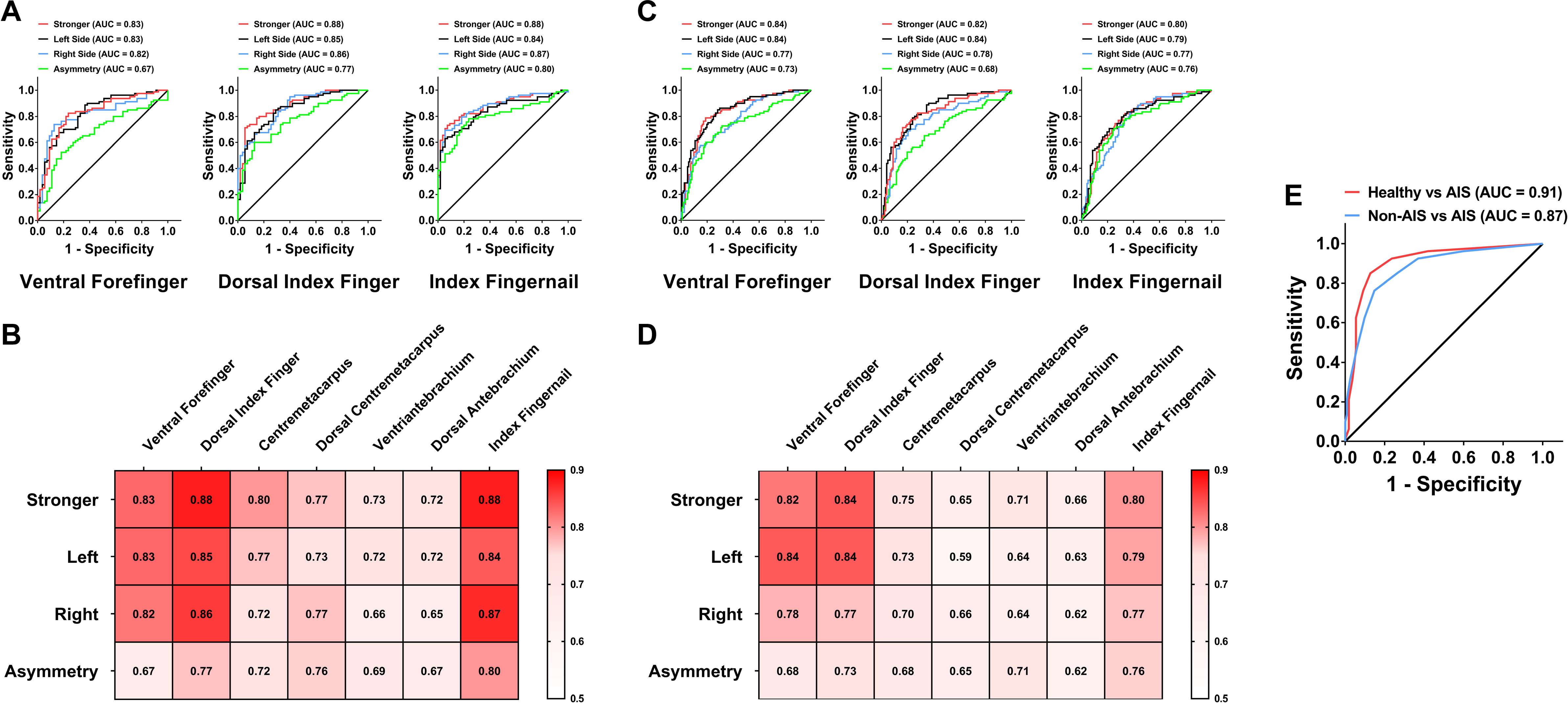
ROC analyses on the AF properties showed significant promise of our AF- based diagnostic method for AIS. (A) By using the AF intensity of the Stronger Side of the Dorsal Index Fingers, the Index Fingernails and the Ventral Forefingers, the AUC for differentiating the AIS patients from the healthy subjects was 0.88, 0.88 and 0.83, respectively. (B) By using the AF intensity at the Stronger Side, the left side or the right side of each examined position or using the AF asymmetry as the variable, ROC analyses were conducted to determine the AUCs for differentiating the AIS patients from the healthy subjects. (C) By using the AF intensity of the Stronger Side of the Dorsal Index Fingers, the Stronger Side of the Index Fingernails, or the left side of the Ventral Forefingers as the variable, the AUC for differentiating the AIS patients from the Non-AIS subjects was 0.84, 0.80 and 0.84, respectively. (D) By using the AF intensity at the Stronger Side, the left side or the right side of each examined position or using the AF asymmetry as the variable, ROC analyses were conducted to determine the AUCs for differentiating the AIS patients from the Non-AIS subjects. (E) By using the number of the examined positions with increased AF intensity as the variable, ROC analyses were conducted to determine the AUC for differentiating the AIS patients from the healthy subjects as well as the AUC for differentiating the AIS patients from the Non-AIS subjects: The AUC was 0.91 and 0.87, respectively. The number of the healthy subjects, the Non-AIS subjects and the AIS subjects was 55, 122, 80, respectively.

By using the AF intensity at the Stronger Side, the left and or right side of each examined position or using the AF asymmetry as the variable, ROC analyses were also conducted to determine the AUCs for differentiating the AIS patients from the Non-AIS subjects (Fig. 5, C and D). In particular, by using the AF intensity of the Stronger Side of the Dorsal Index Fingers, the Stronger Side of the Index Fingernails, or the left Ventral Forefingers as the variable, the AUC for differentiating the AIS patients from the Non-AIS subjects was 0.84, 0.80 and 0.84, respectively (Fig. 5, C and D).

By using the number of the examined positions with increased AF intensity as the variable, ROC analyses were also conducted to determine the AUC for differentiating the AIS patients from the healthy subjects as well as the AUC for differentiating the AIS patients from the Non-AIS subjects: The AUC was 0.91 and 0.87, respectively (Fig. 5E). The specificity and sensitivity for differentiating the AIS patients from the healthy subjects and those for differentiating the AIS patients from the Non-AIS subjects were listed in Table 1.

**Table 1.** The specificity and sensitivity for differentiating the AIS patients from the healthy subjects and those for differentiating the AIS patients from the Non-AIS subjects. The number of the healthy subjects, the Non-AIS subjects and the AIS subjects was 50, 122, 80, respectively.

| AIS Patients |  |  |  |  |
| --- | --- | --- | --- | --- |
|  | vs |  | vs |  |
|  | Healthy |  | Non-AIS |  |
|  | Sensitivity | Specificity | Sensitivity | Specificity |
| Number of Increased AF |  |  |  |  |
|  | 0.85 | 0.87 | 0.76 | 0.85 |
| Ventral Forefinger |  |  |  |  |
| AF of Stronger Side | 0.80 | 0.78 | 0.78 | 0.77 |
| AF of Left Side | 0.71 | 0.71 | 0.79 | 0.76 |
| AF of Right Side | 0.74 | 0.87 | 0.74 | 0.73 |
| AF Asymmetry | 0.64 | 0.67 | 0.65 | 0.67 |
| Dorsal Index Finger |  |  |  |  |
|  | Sensitivity | Specificity | Sensitivity | Specificity |
| AF of Stronger Side | 0.80 | 0.84 | 0.79 | 0.81 |
| AF of Left Side | 0.76 | 0.76 | 0.80 | 0.75 |
| AF of Right Side | 0.76 | 0.73 | 0.70 | 0.66 |
| AF Asymmetry | 0.74 | 0.67 | 0.73 | 0.69 |
| Centremetacarpus |  |  |  |  |
| AF of Stronger Side | 0.71 | 0.73 | 0.69 | 0.69 |
| AF of Left Side | 0.74 | 0.73 | 0.70 | 0.67 |
| AF of Right Side | 0.66 | 0.65 | 0.61 | 0.63 |
| AF Asymmetry | 0.68 | 0.65 | 0.60 | 0.63 |
| Dorsal Centremetacarpus |  |  |  |  |
| AF of Stronger Side | 0.72 | 0.73 | 0.63 | 0.64 |
| AF of Left Side | 0.70 | 0.71 | 0.63 | 0.56 |
| AF of Right Side | 0.71 | 0.71 | 0.64 | 0.65 |
| AF Asymmetry | 0.69 | 0.73 | 0.61 | 0.63 |
| Ventriantebrachium |  |  |  |  |
| AF of Stronger Side | 0.70 | 0.75 | 0.67 | 0.67 |
| AF of Left Side | 0.72 | 0.67 | 0.60 | 0.59 |
| AF of Right Side | 0.67 | 0.64 | 0.62 | 0.61 |
| AF Asymmetry | 0.71 | 0.65 | 0.64 | 0.66 |
| Dorsal Antebrachium |  |  |  |  |
| AF of Stronger Side | 0.68 | 0.69 | 0.66 | 0.62 |
| AF of Left Side | 0.67 | 0.73 | 0.61 | 0.61 |
| AF of Right Side | 0.68 | 0.62 | 0.62 | 0.62 |
| AF Asymmetry | 0.66 | 0.60 | 0.59 | 0.61 |
| Index Fingernail |  |  |  |  |
| AF of Stronger Side | 0.81 | 0.82 | 0.74 | 0.74 |
| AF of Left Side | 0.76 | 0.75 | 0.74 | 0.73 |
| AF of Right Side | 0.82 | 0.78 | 0.72 | 0.71 |
| AF Asymmetry | 0.78 | 0.76 | 0.72 | 0.73 |

By analyzing the AF images of each examined position, we also conducted machine learning-based analyses on the AUCs for differentiating the AIS patients from the healthy subjects (Table 2): By analyzing the AF images of the Dorsal Index Fingers, the Index Fingernails, or the Dorsal Centremetacarpus, the AUC for differentiating the AIS patients from the healthy subjects was 0.91, 0.88, and 0.85, respectively. By analyzing the AF images of each examined position, we further conducted machine learning-based analyses on the AUCs for differentiating the AIS patients from the Non-AIS subjects (Table 2): By analyzing the skin’s AF images of the Dorsal Index Fingers or the Ventral Forefingers, the AUC for differentiating the AIS patients from the Non-AIS subjects was 0.84 and 0.81, respectively.

**Table 2.**
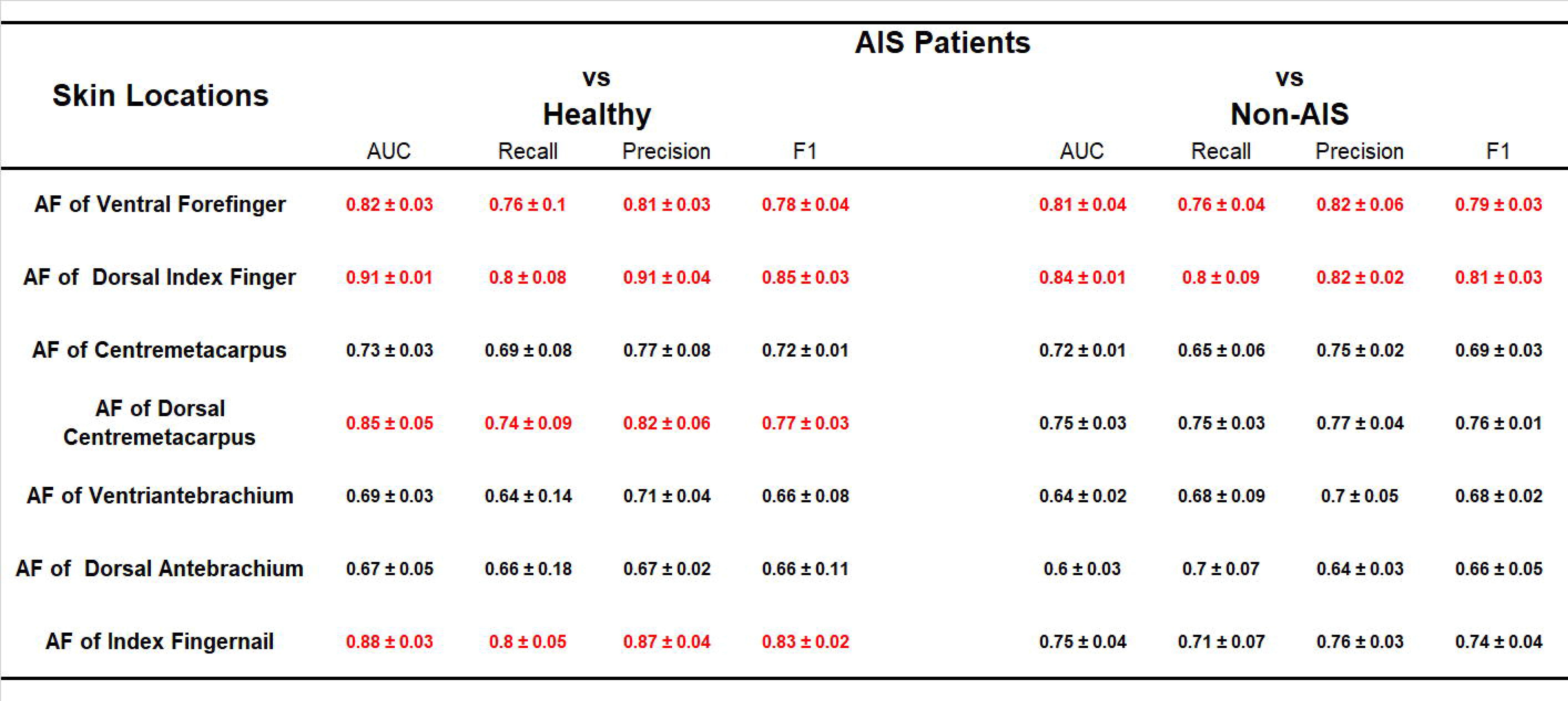
Machine learning-based analyses on the AUCs for differentiating the AIS patients from the healthy subjects and for differentiating the AIS patients from the Non-AIS subjects. By analyzing the AF images of each examined position, machine learning-based analyses were conducted to obtain the AUCs for differentiating the AIS patients from the healthy subjects. By analyzing the AF images of each examined position, machine learning-based analyses were also conducted to obtain the AUCs for differentiating the AIS patients from the Non-AIS subjects. The number of the healthy subjects, the Non-AIS subjects and the AIS subjects was 50, 122, 80, respectively.

We further applied machine learning-based approach to search for the optimal combinations of the AF properties of the examined positions so as to achieve optimal AUCs (Table 3): By using the combined information of the AF properties of the Dorsal Index Fingers and those of the Ventral Forefingers, an optimal AUC of 0.93 was achieved for differentiating the AIS patients from the healthy subjects, with the value of Recall, Precision and F1 being 0.84, 0.93 and 0.88, respectively. By analyzing the combined information of the AF properties of the Dorsal Index Fingers and those of the Ventral Forefingers, an optimal AUC of 0.87 was achieved for differentiating the AIS patients from the Non-AIS subjects, with the value of Recall, Precision and F1 being 0.81, 0.84 and 0.83, respectively.

**Table 3.** Machine learning-based approach to search for the optimal combinations of the AF properties for obtaining the optimal AUC for differentiating the AIS patients from the healthy subjects and for differentiating the AIS patients from the Non-AIS subjects. By analyzing the combined information of the AF properties of the Dorsal Index Fingers and those of the Index Fingernails, an optimal AUC of 0.93 was achieved for differentiating the AIS patients from the healthy subjects. By analyzing the combined information of the AF properties of the Dorsal Index Fingers and those of the Ventral Forefingers, an optimal AUC of 0.87 was achieved for differentiating the AIS patients from the Non-AIS subjects. The number of the healthy subjects, the Non-AIS subjects and the AIS subjects was 50, 122 and 80, respectively.

|  | AIS Patients |  |
| --- | --- | --- |
|  | vs<br>Healthy | vs<br>Non-AIS |
| Skin Locations | Dorsal Index Finger +<br>Index Fingernail | Dorsal Index Finger +<br>Ventral Forefinger |
| AUC | 0.93 ± 0.04 | 0.87 ± 0.02 |
| Recall | 0.84 ± 0.02 | 0.81 ± 0.02 |
| Precision | 0.93 ± 0.05 | 0.84 ± 0.04 |
| F1 | 0.88 ± 0.01 | 0.83 ± 0.01 |

### 5. Logistic regression analyses on the categorical variables, the continuous variables and the AF properties of AIS patients and Non-AIS patients

The categorical variables and the continuous variables of the Non-AIS subjects and the AIS patients were shown in Table S2. In the multivariate logistic regression analyses on the variables and AF Intensity at the Stronger Side, there was statistical significance at all examined positions (Table S3-9). In the multivariate logistic regression analyses on the variables and AF Intensity at the left side, there was statistical significance at all examined positions except Dorsal Centremetacarpus (Table S10-S16).

In the multivariate logistic regression analyses on the variables and AF Intensity at the right side, there was statistical significance at all examined positions except Dorsal Antebrachium (Table S17-S23). In the multivariate logistic regression analyses on the variables and AF asymmetry, there was statistical significance at all examined positions except Dorsal Antebrachium and Index Fingernail (Table S24-S30).

### 6. The AF properties of the AIS patients were significantly different from those of the patients of PD, PI and TIA

We conducted the comparisons among the AF intensity of the AIS patients, the PD patients and the PI patients (Fig. 6, A to G): The AIS patients had significantly higher AF intensity than that of the PD patients only at the Stronger side, the left side and the right side of the Index Fingernails and the skin’s positions of Ventral Forefingers and Dorsal Index Fingers. The AIS patients also had significantly higher AF intensity than that of the PI patients at the Stronger side, the left side and the right side of the Index Fingernails and the skin’s positions of the Stronger side, the left side and the right side of the Dorsal Index Fingers, Ventral Forefingers, and Dorsal Centremetacarpus. Moreover, the skin’s AF intensity of the AIS patients was significantly higher than that of the TIA patients only at the Strong Side, left side and right side of the Index Fingernails and the skin’s position of the Ventral Forefingers as well as the Stronger Side of the Centremetacarpus (Fig. S5, A to G).

**Fig. 6.**
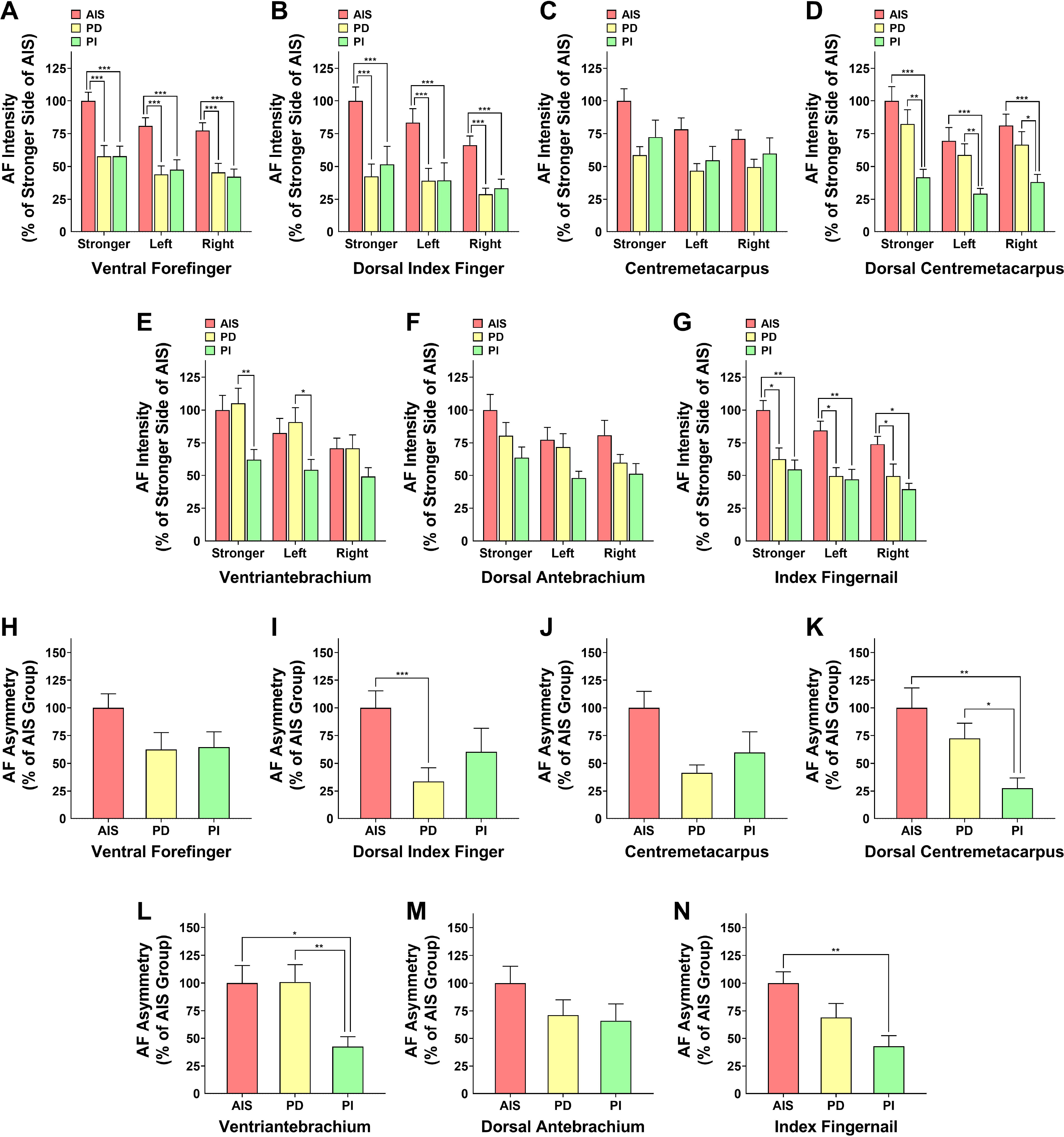
The AF properties of the AIS patients were significantly different from those of the patients of PD and PI. (A-G) The AIS patients had significantly higher AF intensity than that of the PD patients only at the Stronger side, the left side and the right side of the Index Fingernails and the skin’s positions of Ventral Forefingers and Dorsal Index Fingers. The AIS patients also had significantly higher AF intensity than that of the PI patients at the Stronger side, the left side and the right side of the Index Fingernails and the skin’s positions of the Stronger side, the left side and the right side of the Dorsal Index Fingers, Ventral Forefingers, and Dorsal Centremetacarpus. (H-N) The AF asymmetry of the AIS patients was significantly higher than that of the PD patients only at the Dorsal Index Fingers. The AF asymmetry of the AIS patients was also significantly higher than that of the PI patients at the Index Fingernails and the skin’s position of Dorsal Centremetacarpus and Ventriantebrachium. The number of the AIS patients, the PD patients and the PI patients was 67-80, 30 and 29, respectively. *, *P* < 0.05; **, *P* < 0.01; ***, *P* < 0.001.

We also conducted the comparisons among the AF asymmetry of the AIS patients, the PD patients and the PI patients (Fig. 6, H to N): The AF asymmetry of the AIS patients was significantly higher than that of the PD patients only at the Dorsal Index Fingers. The AF asymmetry of the AIS patients was also significantly higher than that of the PI patients at the Index Fingernails and the skin’s position of Dorsal Centremetacarpus and Ventriantebrachium. Moreover, the skin’s AF asymmetry of the AIS patients was significantly higher than that of the TIA patients only at the Index Fingernails and the skin’s positions of Ventral Forefingers (Fig. S6, A to G).

At both right and left Index Fingernails and the skin’s positions of right and left Dorsal Index Fingers and Ventral Forefingers, the AIS patients had remarkably higher percentages of the examined positions with increased AF, compared to those of PD patients (Fig. S7A). At all examined positions except the left Centremetacarpus, the AIS patients had remarkably higher percentages of the examined positions with increased AF, compared to those of the PI patients (Fig. S7A). At the 14 examined positions, the percentage of the PD patients with > 3 positions with increased AF intensity was 33%, while the percentage of the PI patients with > 3 positions with increased AF intensity was 34% (Fig. S7B). In contrast, the percentage of the AIS patients with > 3 positions with increased AF intensity was 76% (Fig. S7B).

### 7. ROC analyses and machine learning-based analyses on the AF properties showed promising AUCs for differentiating AIS patients from the patients of PD, PI, and TIA

By using the AF intensity at the Stronger Side, the left side or the right side of each examined position or using the AF asymmetry as the variable, ROC analyses were conducted to determine the AUCs for differentiating the AIS patients from the PD patients (Fig. 7A) as well as the AUCs for differentiating the AIS patients from the PI patients (Fig. 7B). In particular, by analyzing the skin’s AF intensity of the Stronger Side of the Dorsal Index Fingers or the left Ventral Forefingers, the AUC for differentiating the AIS patients from the PD patients was 0.79 and 0.77, respectively (Fig. 7A). By analyzing the AF intensity of the left Dorsal Index Fingers, the left Dorsal Centremetacarpus or the left Ventral Forefingers as the variable, the AUC for differentiating the AIS patients from the PI patients was 0.80, 0.76 and 0.75, respectively (Fig. 7B).

**Fig. 7.**
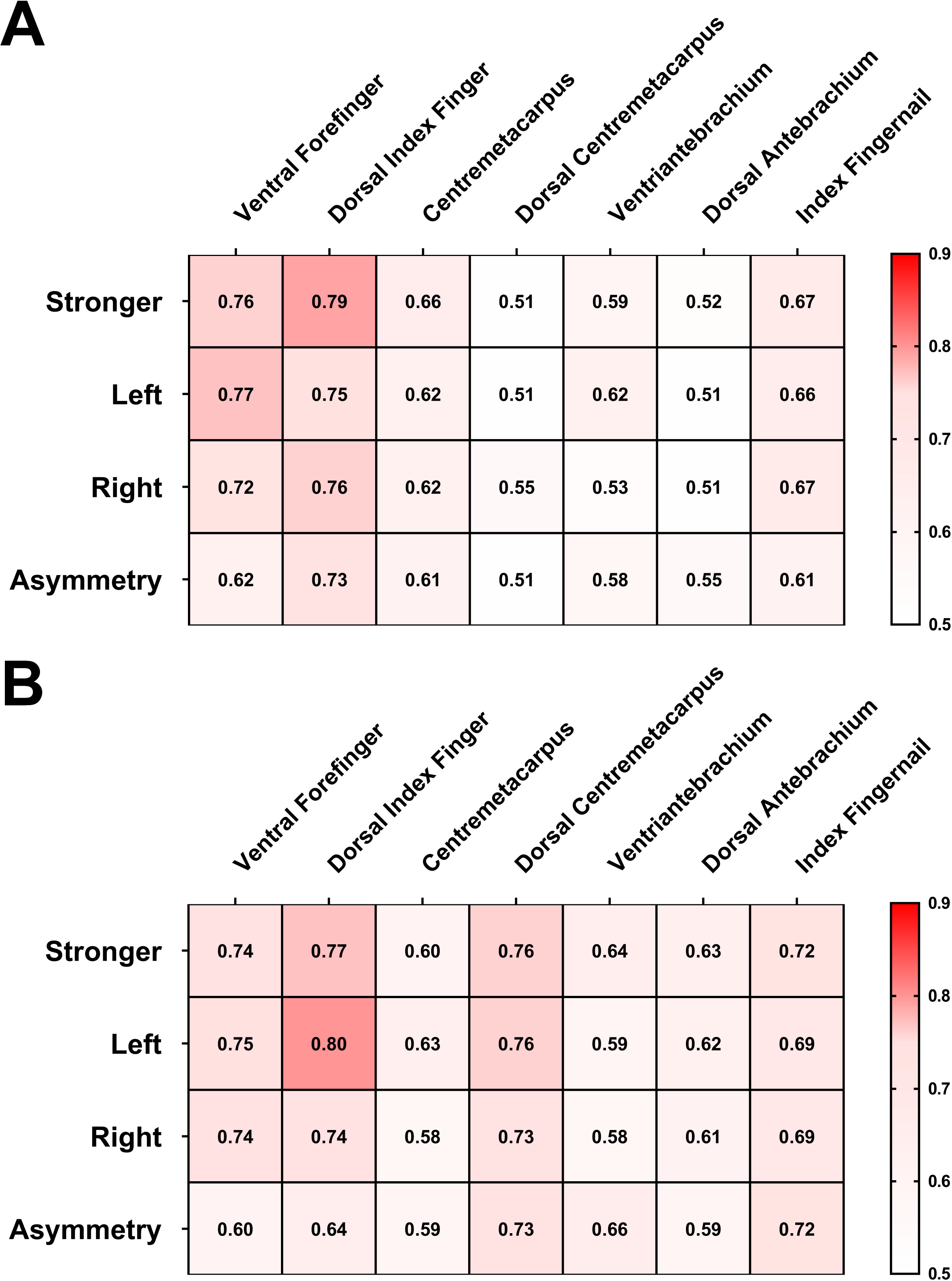
ROC analyses on the AF properties showed promising AUCs for differentiating the AIS patients from the PD patients and for differentiating the AIS patients from the PI patients. (A) By using the AF intensity at the Stronger Side, the left side or the right side of each examined position or using the AF asymmetry as the variable, ROC analyses were conducted to determine the AUCs for differentiating the AIS patients from the PD patients. (B) By using the AF intensity at the Stronger Side, the left and or right side of each examined position or using the AF asymmetry as the variable, ROC analyses were also conducted to determine the AUCs for differentiating the AIS patients from the PI patients. The number of the AIS patients, the PD patients and the PI patients was 67-80, 30 and 29, respectively.

By using the AF intensity at the Stronger Side, the left side or the right side of each examined position or using the AF asymmetry as the variable, ROC analyses were also conducted to determine the AUCs for differentiating the AIS patients from the TIA patients (Fig. S8). In particular, by using the AF intensity of the Stronger Side of the Ventral Forefingers as the variable, the AUC for differentiating the AIS patients from the TIA patients was 0.83 (Fig. S8).

By analyzing the AF images of each examined position of these groups, we further conducted machine learning-based analyses for the AUC for differentiating the AIS patients from the PD patients and the AUC for differentiating the AIS patients from the PI patients (Table 4). In order to achieve optimal AUCs, we further applied machine learning-based approach to search for the optimal combinations of the AF properties of the examined positions (Table 5): By using the combined information of the skin’s AF properties of the Ventral Forefingers and Centremetacarpus, an optimal AUC of 0.79 was achieved for differentiating the AIS patients from the PD patients, with the value of Recall, Precision and F1 being 0.79, 0.83, and 0.80, respectively. By using the combined information of the skin’s AF properties of the Dorsal Index Fingers and the Ventral Forefingers, an optimal AUC of 0.82 was achieved for differentiating the AIS patients from the PI patients, with the value of Recall, Precision and F1 being 0.93, 0.84, and 0.88, respectively.

**Table 4.**
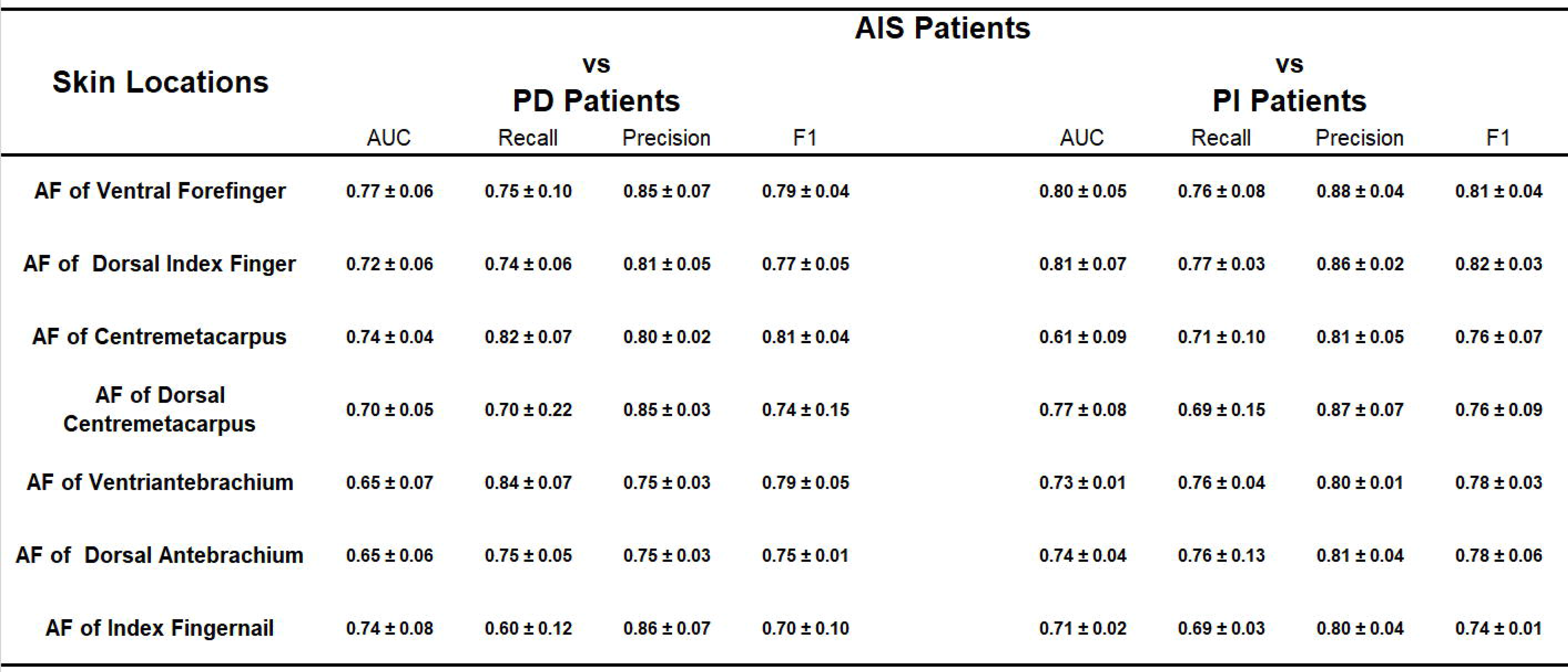
Machine learning-based analyses on the AUC for differentiating the AIS patients from the PD patients and for differentiating the AIS patients from the PI patients. By using the AF properties of each examined position of these groups, machine learning-based analyses were conducted for the AUC for differentiating the AIS patients from the PD patients and the AUC for differentiating the AIS patients from the PI patients. The number of the AIS patients, the PD patients and the PI patients was 80, 30 and 29, respectively.

**Table 5.** Machine learning-based searches for the optimal combinations of the AF properties for obtaining optimal AUCs for differentiating the AIS patients from the PD patients and for differentiating the AIS patients from the PI patients. By analyzing the combined information of the skin’s AF properties of the Ventral Forefingers and Centremetacarpus, an optimal AUC of 0.79 was achieved for differentiating the AIS patients from the PD patients, with the value of Recall, Precision and F1 being 0.79, 0.83, and 0.80, respectively. By analyzing the combined information of the skin’s AF properties of the Dorsal Index Fingers and the Ventral Forefingers, an optimal AUC of 0.82 was achieved for differentiating the AIS patients from the PI patients, with the value of Recall, Precision and F1 being 0.93, 0.84, and 0.88, respectively. The number of the AIS patients, the PD patients and the PI patients was 80, 30, and 29, respectively.

|  | AIS Patients |  |
| --- | --- | --- |
|  | vs<br>PD Patients | vs<br>PI Patients |
| Skin Locations | Ventral Forefinger<br>+<br>Centrometacarpus | Dorsal Index Finger<br>+<br>Ventral Forefinger |
| AUC | 0.79 ± 0.11 | 0.82 ± 0.03 |
| Recall | 0.79 ± 0.09 | 0.93 ± 0.03 |
| Precision | 0.83 ± 0.07 | 0.84 ± 0.05 |
| F1 | 0.80 ± 0.07 | 0.88 ± 0.01 |

By using the AF properties of the Index Fingernails and the skin’s examined positions, machine learning-based analyses on the AUC for differentiating the AIS patients from the TIA patients were also conducted (Table S31). In particular, by using the skin’s AF intensity of the Ventral Forefingers or the Dorsal Centremetacarpus as the variable, the AUC for differentiating the AIS patients from the TIA patients was 0.87 and 0.81, respectively (Table S31). By using the combined information of the skin’s AF properties of the Ventral Forefingers and the Dorsal Centremetacarpus as the variable, an optimal AUC of 0.88 was achieved for differentiating the AIS patients from the TIA patients, with the value of Recall, Precision and F1 being 0.88, 0.94 and 0.90, respectively (Table S32).

### 8. Analyses on the relationships between the AIS patients’ AF properties and NIHSS scores

NIHSS score is an index of neurological functional deficits, which is valuable for early prognostication and serial assessment^19^. We found that the AIS patients with NIHSS scores equal to or higher than 4 had significantly higher AF intensity at the skin’s position of both Strong Side and the left side of Dorsal Index Fingers as well as the left Dorsal Antebrachium, compared to the AIS patients with NIHSS scores equal to or lower than 3 (Fig. 8, A to G). We further conducted machine learning-based analyses on the relationships between the AF images and the NIHSS scores of the AIS patients (Table 6): By analyzing the AF images of the Index Fingernails and the skin’s positions of the Ventral Forefingers, Dorsal Index Fingers, Dorsal Centremetacarpus and Dorsal Antebrachium, the AUC for differentiating the AIS patients with NIHSS scores equal to or higher than 4 from the AIS patients with NIHSS scores equal to or lower than 3 was 0.70, 0.70, 0.70, 0.72, and 0.71, respectively.

**Fig. 8.**
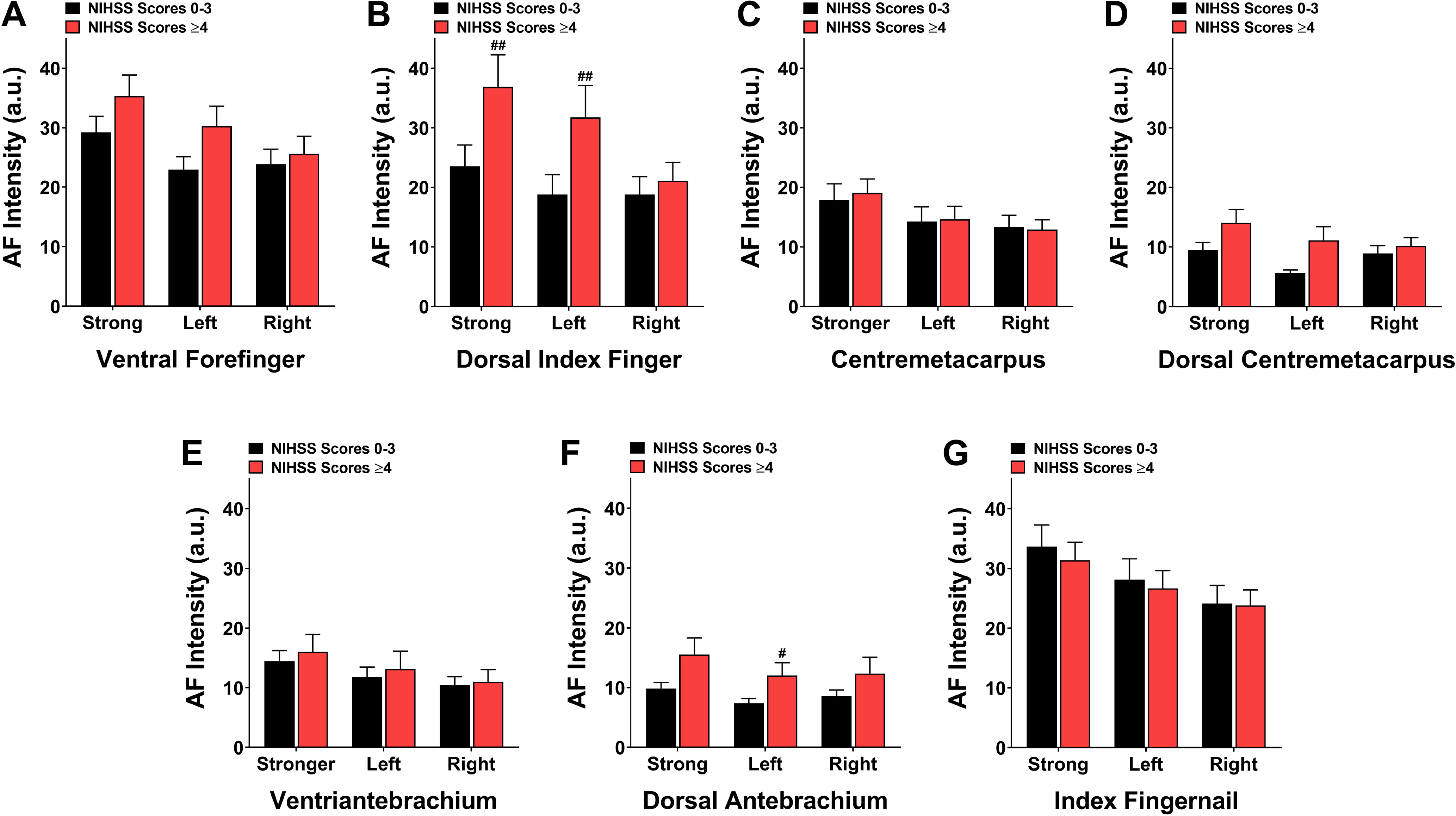
Analyses on the relationships between the AIS patients’ AF properties and NIHSS scores. (A-G) The AIS patients with NIHSS scores equal to or higher than 4 had significantly higher AF intensity at the skin’s position of both Strong Side and the left side of Dorsal Index Fingers as well as the left Dorsal Antebrachium, compared to the AIS patients with NIHSS scores equal to or lower than 3. The number of the AIS patients with NIHSS scores equal to or smaller than 3 and the number of the AIS patients with NIHSS scores equal to or higher than 4 was 38 and 40, respectively. #, *p* < 0.05; ##, *p* < 0.01 (M-W test).

**Table 6.**
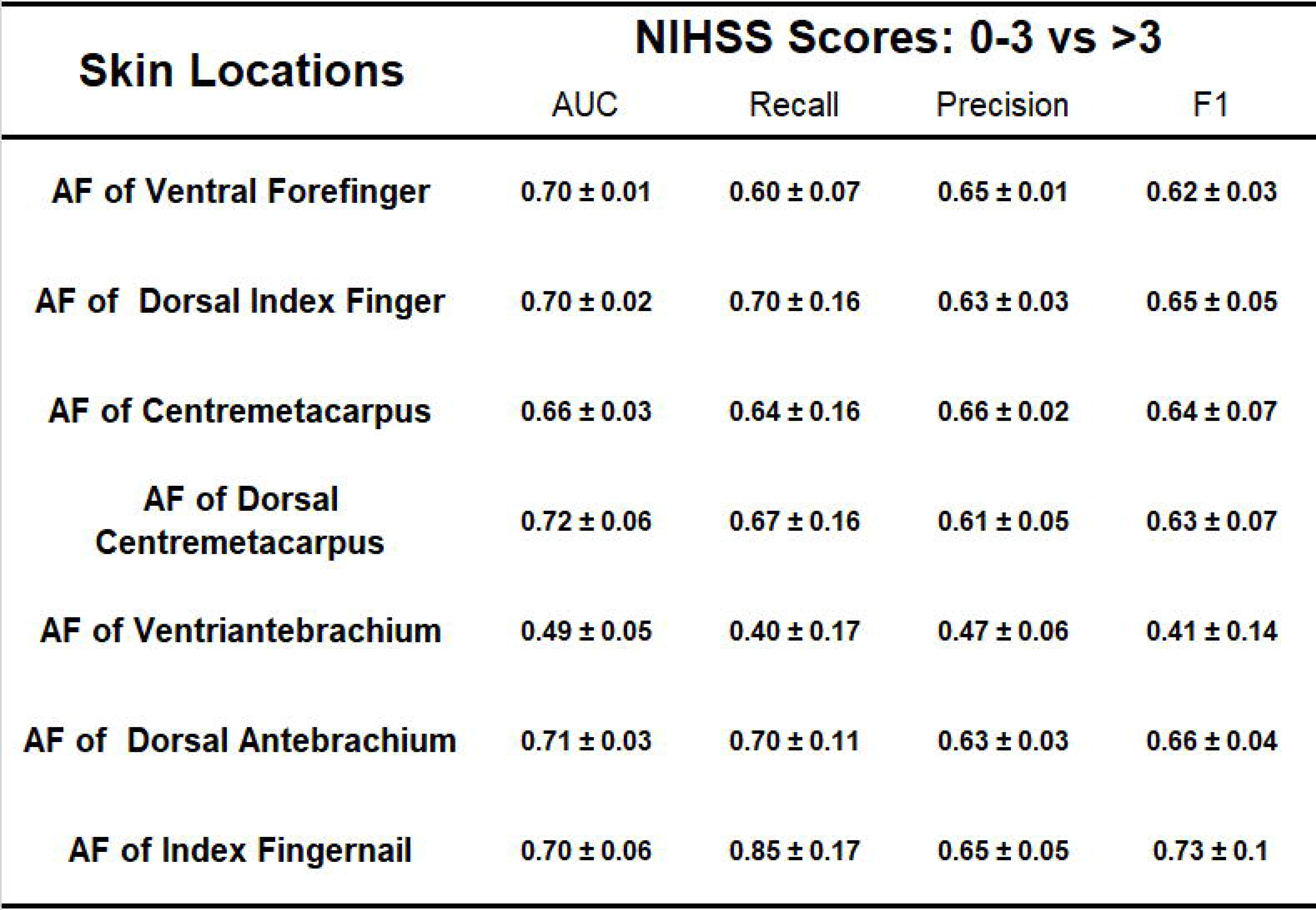
Machine learning-based analyses on the relationships between the AF images and the NIHSS scores of the AIS patients. By analyzing the AF images of the Index Fingernails and the skin’s positions of the Ventral Forefingers, that Dorsal Index Fingers, Dorsal Centremetacarpus and the Dorsal Antebrachium, the AUC for differentiating the AIS patients with NIHSS scores equal to or higher than 4 from the AIS patients with NIHSS scores equal to or lower than 3 was 0.70, 0.70, 0.70, 0.72, and 0.71, respectively. The number of the AIS patients with NIHSS scores equal to or smaller than 3 and the number of the AIS patients with NIHSS scores equal to or higher than 4 was 38 and 40, respectively.

## Discussion

Our study has shown that the AIS patients had significant differences in several AF properties at multiple examined positions, compared to those of all groups examined, including the healthy subjects, the Non-AIS subjects, the PD patients, the PI patients, the TIA patients and the patients of the Recovery Group. The AF properties include AF intensity, AF asymmetry, number of the examined positions with increased AF intensity, and the body’s positions with increased AF. These significant differences in the AF properties have established essential basis for differentiating the AIS patients from the other groups of subjects. As shown by our ROC analyses and machine learning-based analyses on the AF properties, the AUCs have reached highly promising levels for differentiating the AIS patients from the other groups of subjects. In particular, the AUC for differentiating the AIS patients from the healthy subjects and the AUC for differentiating the AIS patients from the Non-AIS subjects reached 0.93 and 0.87, respectively. These findings have strongly indicated that our novel AF-based method holds excellent potential to become a non-invasive, label-free and economical diagnostic approach for AIS. We expect that the AUC for our AF-based AIS diagnosis would be further significantly enhanced, with additions of artificial intelligence (AI)- based analyses on clinical symptoms, increases in the AF data, and AI-based analyses on the AF images.

We proposed a novel concept to define the collective and characteristic properties of these AF properties of AIS patients - the ‘Pattern of AF’ of AIS: The ‘Pattern of AF’ is defined as the combinations of characteristic AF properties at multiple examined positions, which includes several AF properties stated above, as well as the structural properties of the AF images. The AIS’s ‘Pattern of AF’ could become an essential basis for establishing a novel diagnostic technology for AIS. The key implication of the ‘Pattern of AF’ is that several AF properties at multiple examined positions are collectively required for achieving optimal capacity for AIS diagnosis. In addition to the information stated above, the significance of the ‘Pattern of AF’ has been further supported by our following findings: By machine learning-based analyses on the AF properties of more than one examined position, we achieved optimal AUC; and optimal AUCs for differentiating AIS patients from different group of subjects can be achieved by analyses on the AF properties of different examined positions. The ‘Pattern of AF Technology’ may also be used for diagnosis of certain other diseases, e.g., our recent study has reported that the AUC for differentiating lung cancer patients from PI patients was 0.87 ^14^. If future studies could demonstrate that the ‘Pattern of AF’ may become a key concept for developing diagnostic approaches for multiple diseases, ‘Pattern of AF Technology’ may become a novel, general medical imaging technology.

Our study has indicated the ‘Pattern of AF Technology’ has the distinct merits that it is not only non-invasive and label-free, but also economical and efficient. To our knowledge, it is the first diagnostic technology for AIS which possesses all the necessary properties for realizing AIS diagnosis or at least auxiliary AIS diagnosis before the patients enter clinical settings. Potential applications of our technology for AIS diagnosis under non-clinical settings may revolutionize AIS diagnosis, which may enable significantly higher number of AIS patients to get early diagnosis. Since our AF-based imaging is markedly more economical compared to MRI and CT imaging, another important potential application of our technology is for AIS diagnosis under settings that do not have sufficient MRI and CT imaging resource.

Our study found that the AIS patients have significantly greater AF asymmetry in the skin and fingernails than that of the other groups, which could be the first report regarding the asymmetrical properties of any biophotonic properties of human body. This AF asymmetry might be accounted for by the following mechanism: The pathological changes on one side of the brains of the AIS patients may cause significant pathological alterations on another side of the patients’ body. However, our analyses on the relationships between the AIS patients’ MRI images and the locations of the blood clots did not show any correlations (data not shown). We proposed another potential mechanism underlying the AF asymmetry: Before the incidence of AIS, the patients had pathological alterations on one side of their body, which led to the AF increases on those locations. Our finding that the High-Risk subjects had significant AF asymmetry at certain examined positions has suggested that the AF asymmetry of the AIS patients could result at least partially from their pre-existing conditions. It is warranted to further investigate the mechanisms underlying the AF asymmetry, which may expose crucial pathogenic mechanisms of AIS.

We found that the AF spectra of all the examined positions of the healthy subjects, the Non-AIS subjects and the AIS patients were highly similar with each other, which match that of keratins or FAD^17^. Keratins are well-established fluorophores ^21, 22^, which are the main components of human nails ^23^: Human nail plate is composed of approximately 10% - 20% soft epithelial keratins, with the remainder being hard α- keratin ^24^. There is evidence indicating that the increased AF of both the skin and the fingernails of the AIS patients originates from keratins: First, keratins are the sole autofluorescent molecules in the fingernails; and second, the sole autofluorescent molecules that exist in both the skin and the fingernails are keratins. Future studies are necessary to further investigate the AF’s origins.

Our study has found that the Index Fingernails of the AIS patients showed significant and asymmetric increases in the AF, compared to the healthy subjects and the Non-AIS patients. Since human fingernails grow at the rate of approximately 0.1 –0.2 mm / day^20^, the growth of a great majority of the Index Fingernails of the AIS patients had occurred well before our AF determinations. It is reasonable to propose that the asymmetric increases in the Index Fingernails’ AF should occur before the incidence of AIS, since the AIS patients had stroke only 1 - 7 days ago. Therefore, we propose that the ‘Asymmetric Increase in Index Fingernails’ AF’ may become a novel predicative biomarker for AIS incidence.

It is of importance to elucidate the mechanisms underlying the AF increases of the AIS patients. AIS-induced inflammatory responses may play a significant role in the AF increases: Systemic inflammation is one of the major AIS-induced systemic pathological changes^25, 26^, which is a prominent factor that can produce skin’s alterations^27, 28^. Our preliminary study has also indicated that the skin’s green AF intensity is associated with lipopolysaccharide (LPS) dosages in the animal model of inflammation induced by LPS – an inducer of systemic inflammation^29^ (data not shown). Future studies are warranted to further investigate the mechanisms underlying the increased AF of AIS patients.

## Acknowledgment

The authors would like to acknowledge the financial support by two research grants from the Major Special Program Grant of Shanghai Municipality (Grant # 2017SHZDZX01) (to W.Y.), a Major Research Grant from the Scientific Committee of Shanghai Municipality #16JC1400502 (to W.Y.), a Grant for External Applicants, funded by the Fundamental Science & Technology Facility of National Translational Medicine (Shanghai) (Funding # TMSK-2020-133), the Leading Discipline Grant of the Health and Family Planning Commission of Minhang District of Shanghai (#2020MWDXK01) (to D. W.), the Development Grant for Key Specialty of Shanghai Fifth People’s Hospital (#2020WYZDZK04) (to D. W.), and a grant of the Major Research Program of Jiangxi Province (#20212BBG71013).

## Legends of Supplemental Figures

Figure S1. The AF intensity of the High-Risk subjects was significantly higher than that of the healthy subjects at certain examined positions. (A - G) The AF intensity of the High-Risk subjects was significantly higher than that of the healthy subjects only at right Index Fingernails, the skin of Stronger Side and the right side of Dorsal Index Fingers, and the Stronger Side, the left side and the right side of Dorsal Centremetacarpus. The AF intensity of the High-Risk subjects was significantly lower than that of the AIS patients at all examined positions, except at the skin of the Stronger Side, the left side and the right side of Dorsal Centremetacarpus, the skin of the left and the right Antebrachium, and the skin of the right Ventriantebrachium. The number of the healthy subjects, the High-Risk subjects, and the AIS subjects was 55, 59, 78-80, respectively. *, *p* < 0.05; **, *p* < 0.01; ***, *p* < 0.001.

Figure S2. The AF intensity of the patients of the Recovery Group was significantly lower than that of the AIS patients at certain examined positions. (A - G) The subjects of the Recovery Group had significantly lower AF intensity at all examined skin’s positions and the Index Fingernails than that of AIS patients, except at the skin’s positions of the left and right Centremetacarpus, right Ventriantebrachium, right Dorsal Antebrachium, and the Stronger Side and the left side of Dorsal Centremetacarpus. The number of the AIS patients and the patients of the Recovery Group was 78-80 and 37, respectively. #, *p* < 0.05; ##, *p* < 0.01; ###, *p* < 0.001.

Figure S3. The AF asymmetry of the High-Risk subjects was significantly higher than that of the healthy subjects at the skin’s position of Dorsal Centremetacarpus. (A - G) The AF asymmetry of the High-Risk subjects was not significantly different from that of the healthy subjects at all examined positions, except at the skin’s position of Dorsal Centremetacarpus. The AF asymmetry of the High-Risk subjects was significantly lower than that of the AIS patients at all examined positions, except at the skin’s positions of Centremetacarpus and Dorsal Antebrachium. The number of the healthy subjects, the High-Risk subjects and the AIS patients was 55, 59 and 67-80, respectively. *, *p* < 0.05; **, *p* < 0.01; ***, *p* < 0.001.

Figure S4. The High-Risk subjects had higher percentages with increased AF than the healthy subjects at certain examined positions. (A) Higher percentages of the High-Risk subjects had increased AF at certain examined positions, compared to those of the healthy subjects. (B) The percentage of the High-Risk subjects with 2-3, 4-5, or > 5 positions with increased AF intensity was 24%, 4%, 4%, respectively. The number of the healthy subjects, the High-Risk subjects and the AIS patients was 55, 59 and 80, respectively.

Figure S5. TIA patients had significantly lower AF intensity than the AIS patients at certain examined skin’s positions. (A - G) The AF intensity of the AIS patients was significantly higher than that of the TIA patients only at the Stronger Side, the left side, and the right side of the Index Fingernails, the skin’s positions of the Stronger Side, the left side, and the right side of Ventral Forefingers, and the skin’s positions of the Stronger Side of the Centremetacarpus. The number of the AIS patients and the TIA patients was 78-80 and 13, respectively. #, *p* < 0.05; ##, *p* < 0.01; ###, *p* < 0.001.

Figure S6. Comparisons between the AF asymmetry of the AIS patients and that of the TIA patients at examined positions. (A - G) The AF asymmetry of the AIS patients was significantly higher than that of the TIA patients only at the Index Fingernails and the skin’s position of Ventral Forefingers. The number of the AIS patients and the TIA patients was 67-80 and 13, respectively. #, *p* < 0.05; ##, *p* < 0.01.

Figure S7. The AIS patients had remarkably higher percentages of the examined positions with increased AF, compared to those of the PD patients and the PI patients. (A) At both right and left Index Fingernails and the skin’s positions of right and left Dorsal Index Fingers and Ventral Forefingers, the AIS patients had remarkably higher percentages of the examined positions with increased AF, compared to that of PD patients. At all examined positions except the left Centremetacarpus, the AIS patients had remarkably higher percentages of the examined positions with increased AF, compared to that of the PI patients. (B) At the 14 examined positions, the percentage of the PD patients with > 3 positions with increased AF intensity was 33%, while the percentage of the PI patients with > 3 positions with increased AF intensity was 34%. In contrast, the percentage of the AIS patients with > 3 positions with increased AF intensity was 76%. The number of the AIS patients, the PD patients and the PI patients was 80, 30 and 29, respectively.

Figure S8. ROC analyses on the AF properties for obtaining the AUC for differentiating the AIS patients from the TIA patients. By using the AF intensity at the Stronger Side, the left side or the right side of each examined position or using the AF asymmetry as the variable, ROC analyses were conducted to determine the AUCs for differentiating the AIS patients from the TIA patients. The number of the AIS patients and the TIA patients was 67-80 and 13, respectively.

## Legends of Supplemental Tables

Table S1. The baseline information of each group of subjects.

Table S2. The categorical variables and the continuous variables of the Non-AIS subjects and the AIS patients were shown in the table.

Table S3. Multivariate logistic regression analyses on the variables and skin’s AF Intensity at the Stronger Side of Ventral Forefingers.

Table S4. Multivariate logistic regression analyses on the variables and skin’s AF Intensity at the Stronger Side of Dorsal Index Fingers.

Table S5. Multivariate logistic regression analyses on the variables and skin’s AF Intensity at the Stronger Side of Centremetacarpus.

Table S6. Multivariate logistic regression analyses on the variables and skin’s AF Intensity at the Stronger Side of Dorsal Centremetacarpus.

Table S7. Multivariate logistic regression analyses on the variables and skin’s AF Intensity at the Stronger Side of Ventriantebrachium.

Table S8. Multivariate logistic regression analyses on the variables and skin’s AF Intensity at the Stronger Side of Dorsal Antebrachium.

Table S9. Multivariate logistic regression analyses on the variables and the AF Intensity at the Stronger Side of Index Fingernails.

Table S10. Multivariate logistic regression analyses on the variables and skin’s AF Intensity at the left side of Ventral Forefingers.

Table S11. Multivariate logistic regression analyses on the variables and skin’s AF Intensity at the left side of Dorsal Index Fingers.

Table S12. Multivariate logistic regression analyses on the variables and skin’s AF Intensity at the left side of Centremetacarpus.

Table S13. Multivariate logistic regression analyses on the variables and skin’s AF Intensity at the left side of Dorsal Centremetacarpus.

Table S14. Multivariate logistic regression analyses on the variables and skin’s AF Intensity at the left side of Ventriantebrachium.

Table S15. Multivariate logistic regression analyses on the variables and skin’s AF Intensity at the left side of Dorsal Antebrachium.

Table S16. Multivariate logistic regression analyses on the variables and the AF Intensity at the left side of Index Fingernails.

Table S17. Multivariate logistic regression analyses on the variables and skin’s AF Intensity at the right side of Ventral Forefingers.

Table S18. Multivariate logistic regression analyses on the variables and skin’s AF Intensity at the right side of Dorsal Index Fingers.

Table S19. Multivariate logistic regression analyses on the variables and skin’s AF Intensity at the right side of Centremetacarpus.

Table S20. Multivariate logistic regression analyses on the variables and skin’s AF Intensity at the right side of Dorsal Centremetacarpus.

Table S21. Multivariate logistic regression analyses on the variables and skin’s AF Intensity at the right side of Ventriantebrachium.

Table S22. Multivariate logistic regression analyses on the variables and skin’s AF Intensity at the right side of Dorsal Antebrachium.

Table S23. Multivariate logistic regression analyses on the variables and the AF Intensity at the right side of Index Fingernails.

Table S24. Multivariate logistic regression analyses on the variables and skin’s AF asymmetry at the Ventral Forefingers.

Table S25. Multivariate logistic regression analyses on the variables and skin’s AF asymmetry at the Dorsal Index Fingers.

Table S26. Multivariate logistic regression analyses on the variables and skin’s AF asymmetry at the Centremetacarpus.

Table S27. Multivariate logistic regression analyses on the variables and skin’s AF asymmetry at the Dorsal Centremetacarpus.

Table S28. Multivariate logistic regression analyses on the variables and skin’s AF asymmetry at the Ventriantebrachium.

Table S29. Multivariate logistic regression analyses on the variables and skin’s AF asymmetry at the Dorsal Antebrachium.

Table S30. Multivariate logistic regression analyses on the variables and the AF asymmetry at the Index Fingernails.

Table S31. Machine learning-based analyses on the AF properties for obtaining the AUC for differentiating the AIS patients from the TIA patients. By using the AF properties of the Index Fingernails and the skin’s examined positions, machine learning-based analyses on the AUC for differentiating the AIS patients from the TIA patients were conducted.

Table S32. Machine learning-based analyses on the combined information of the AF properties for the optimal AUC for differentiating the AIS patients from the TIA patients. By analyzing the AF images of the Ventral Forefingers and those of the Index Fingernails, an optimal AUC of 0.88 was achieved for differentiating the AIS patients from the TIA patients.

