## Supplementary figures and images for "Distinct ‘pattern of autofluorescence’ of acute ischemic stroke patients’ skin and fingernails: A novel diagnostic biomarker for acute ischemic stroke"

### Supplemental Fig. 1

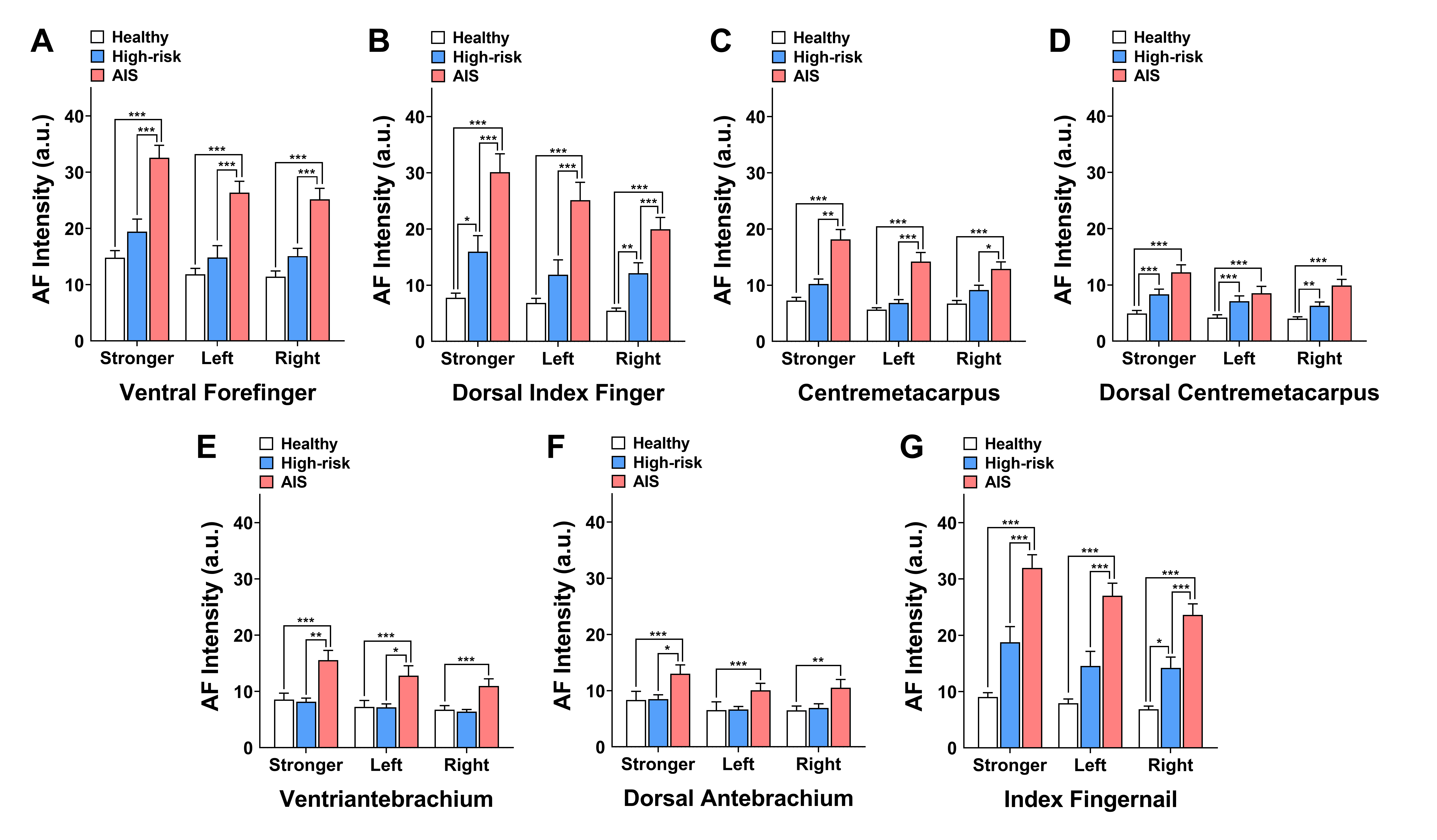

### Supplemental Fig. 2

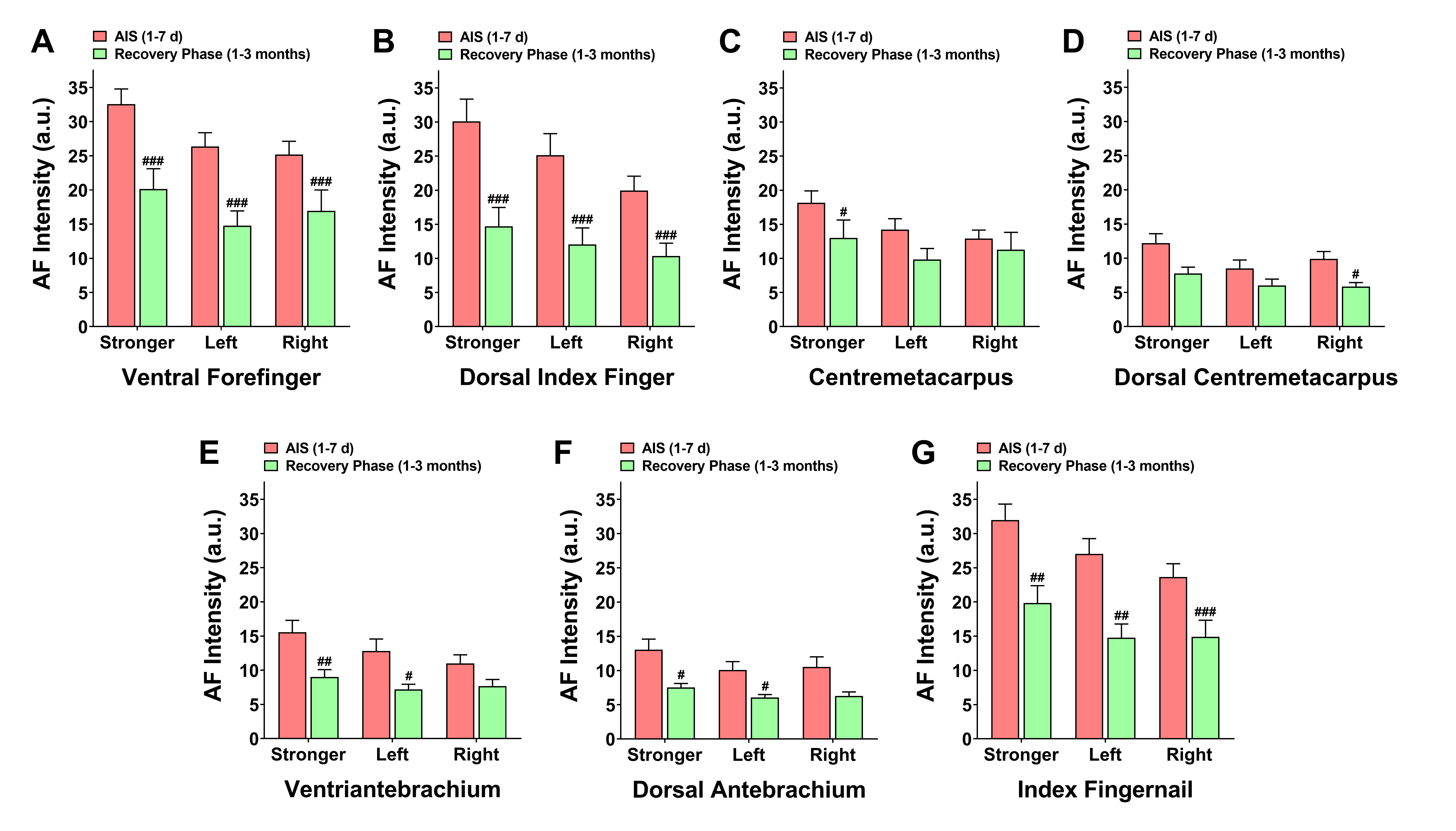

### Supplemental Fig. 3

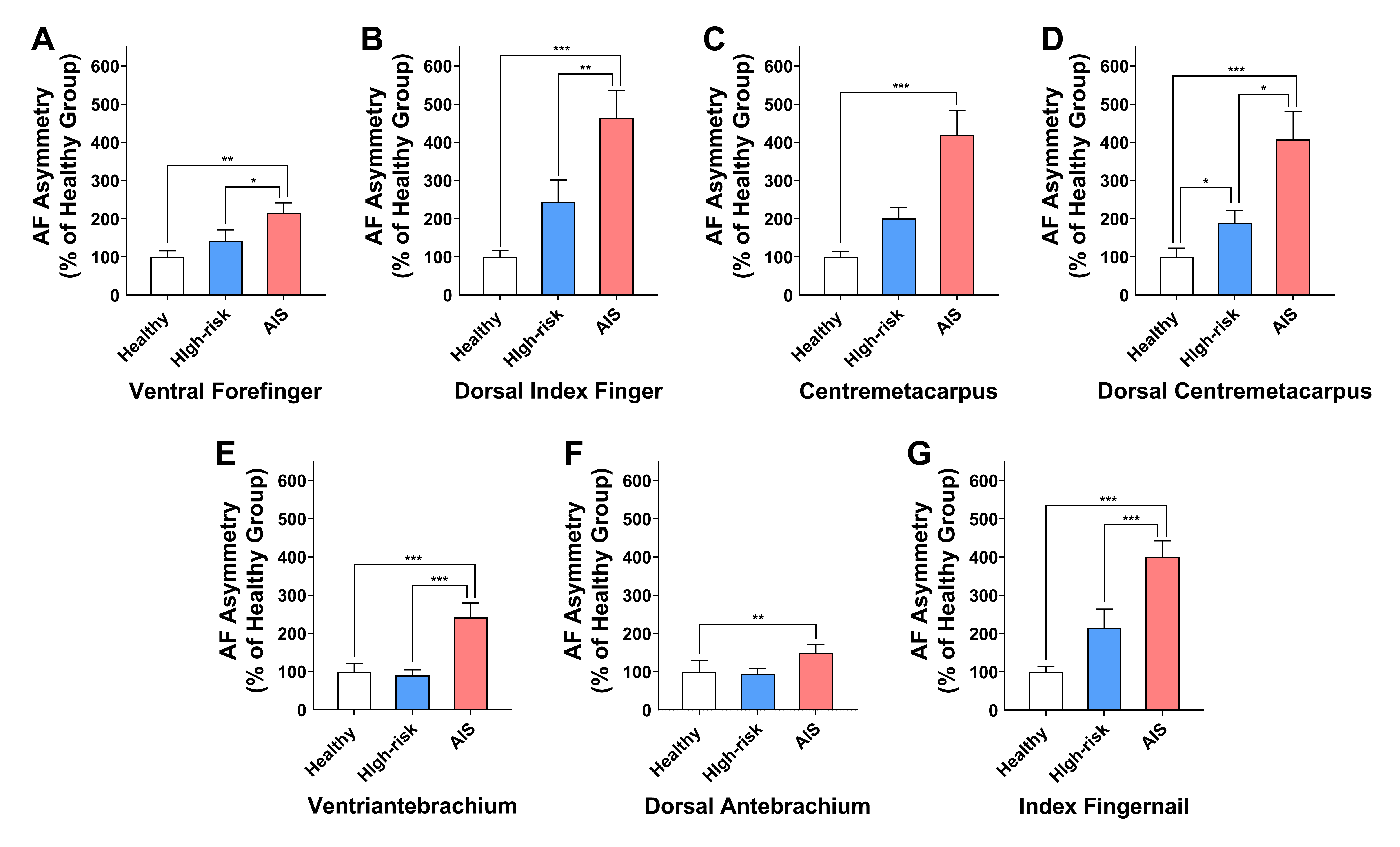

### Supplemental Fig. 4

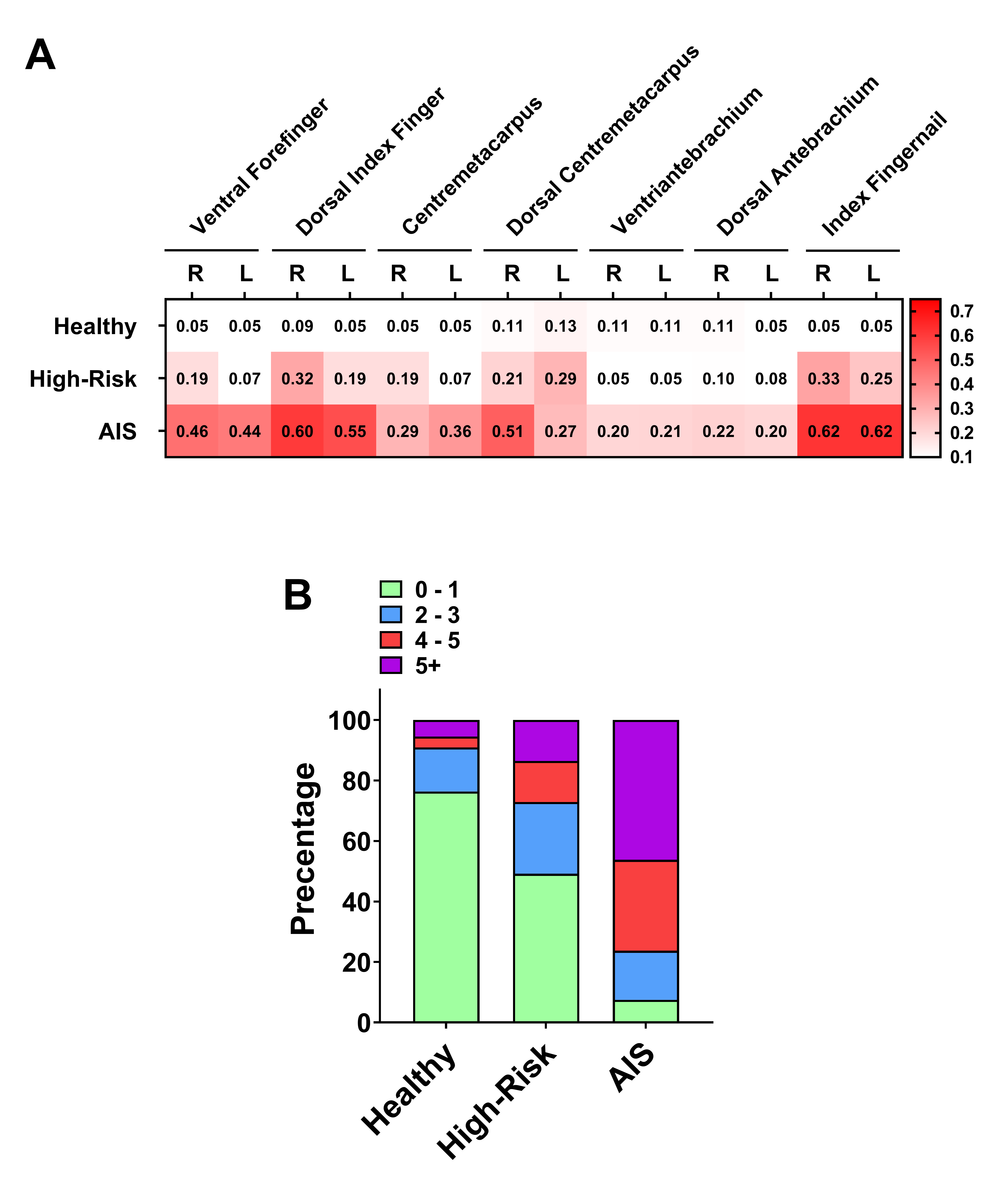

### Supplemental Fig. 5

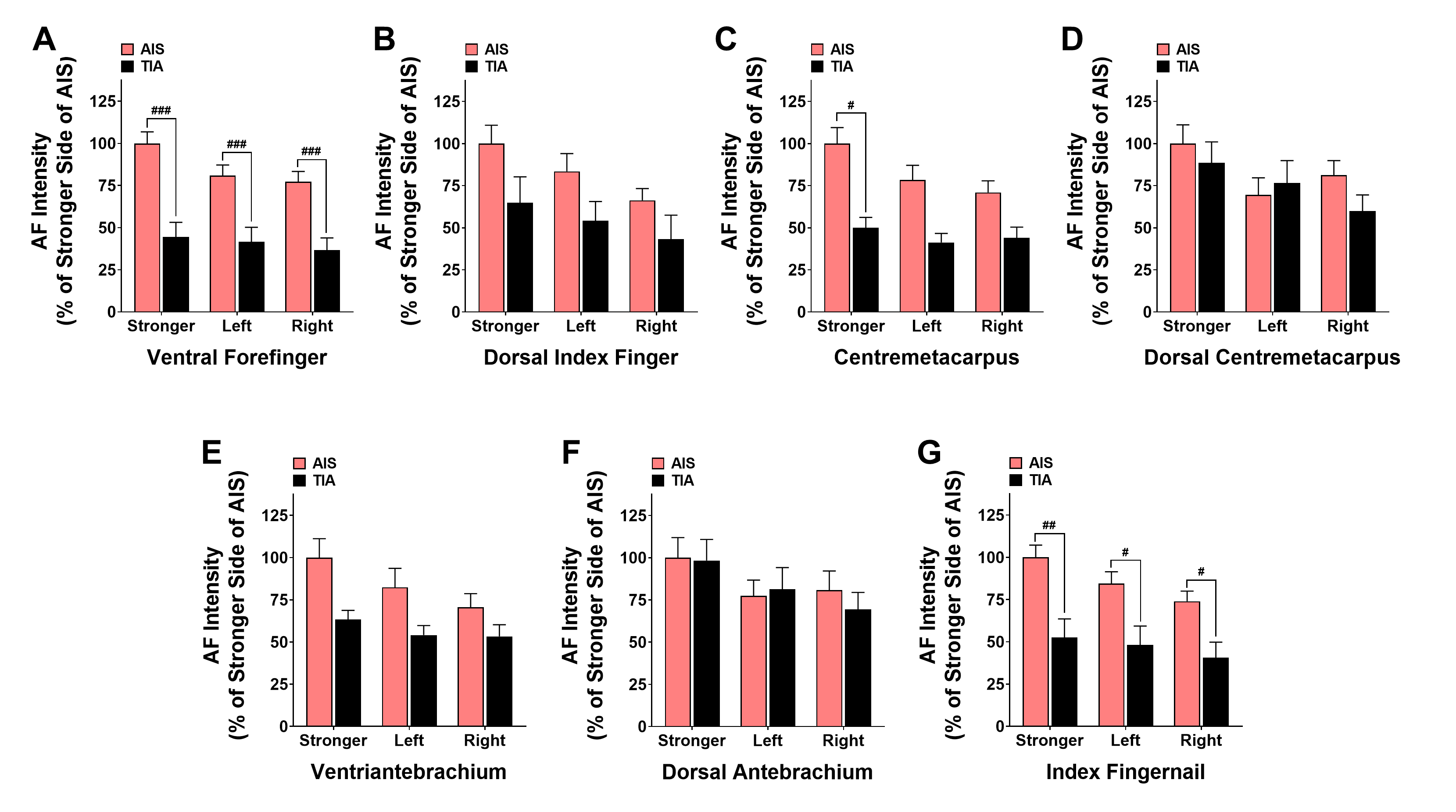

### Supplemental Fig. 6

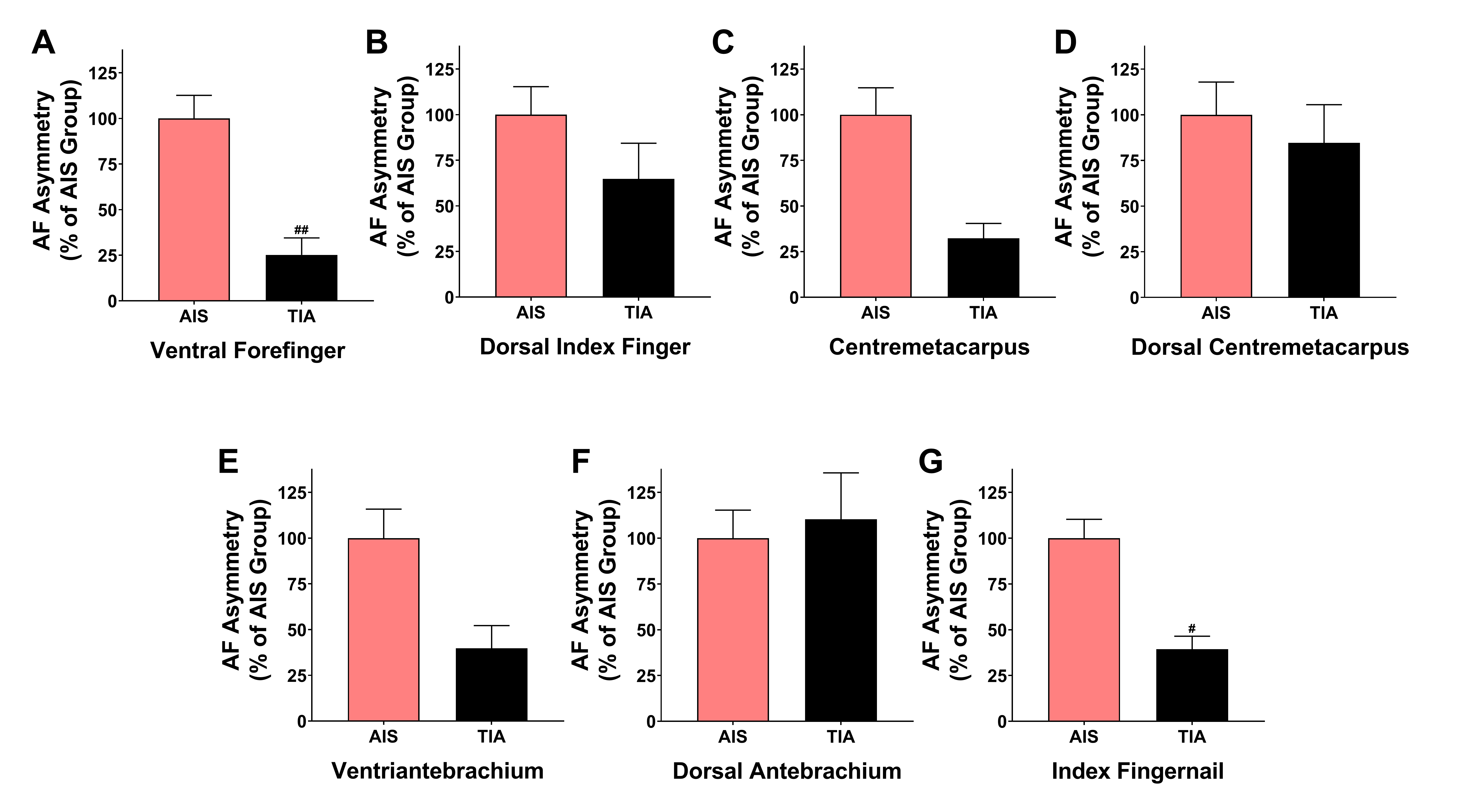

### Supplemental Fig. 7

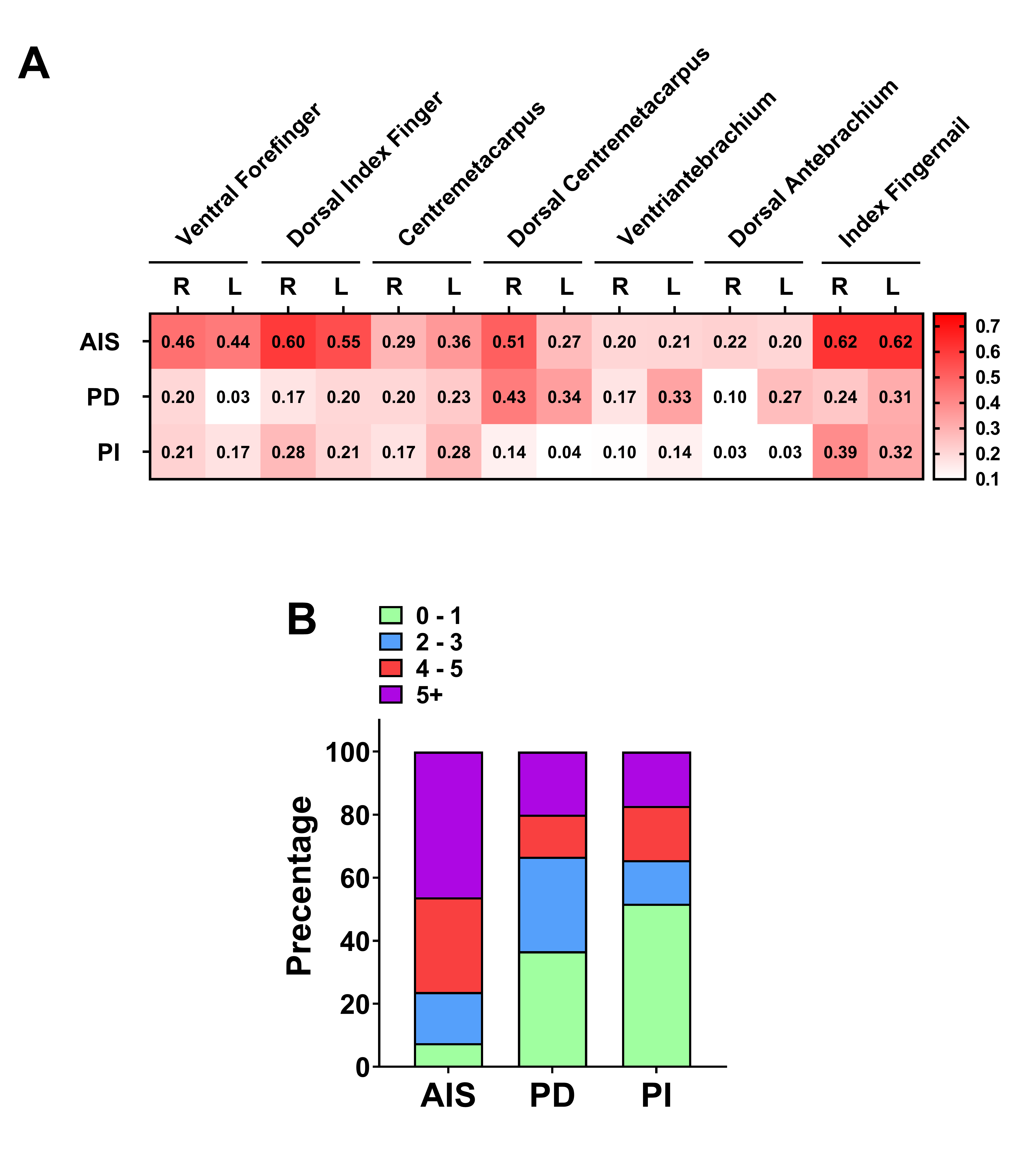

### Supplemental Fig. 8

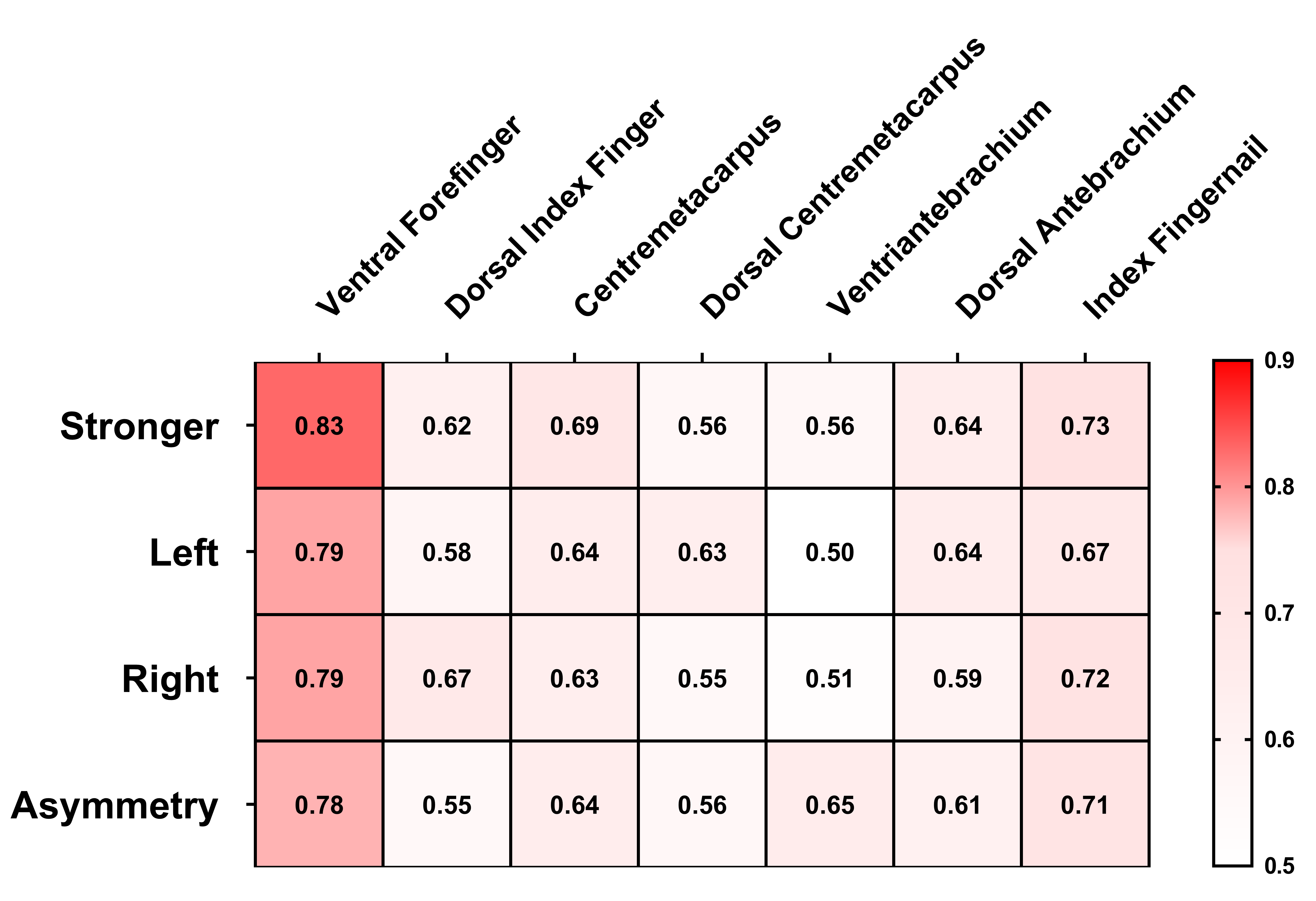

### Supplemental Table 1

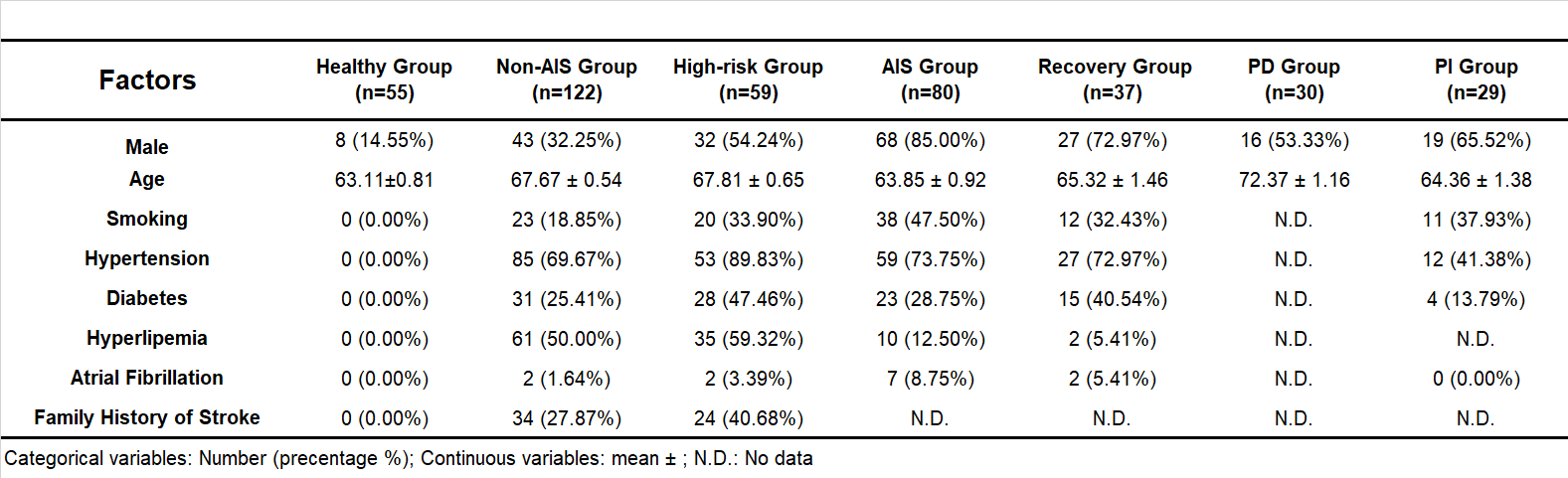

### Supplemental Table 2

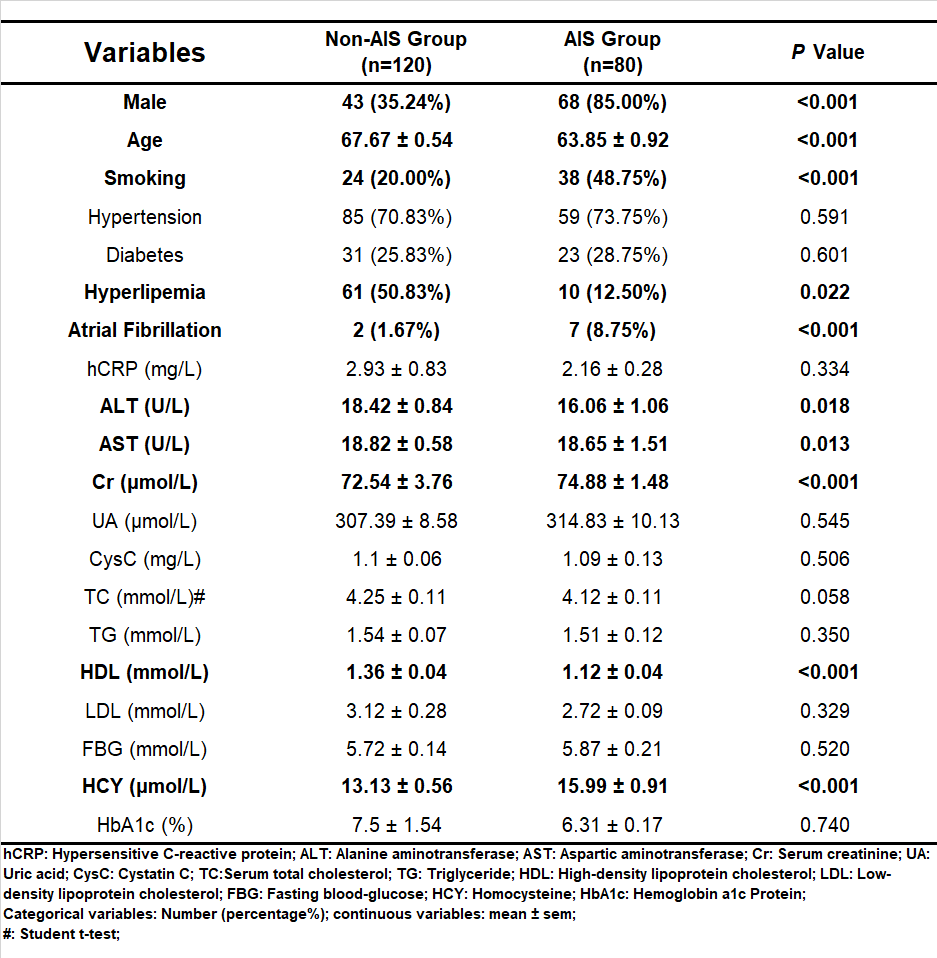

### Supplemental Table 3

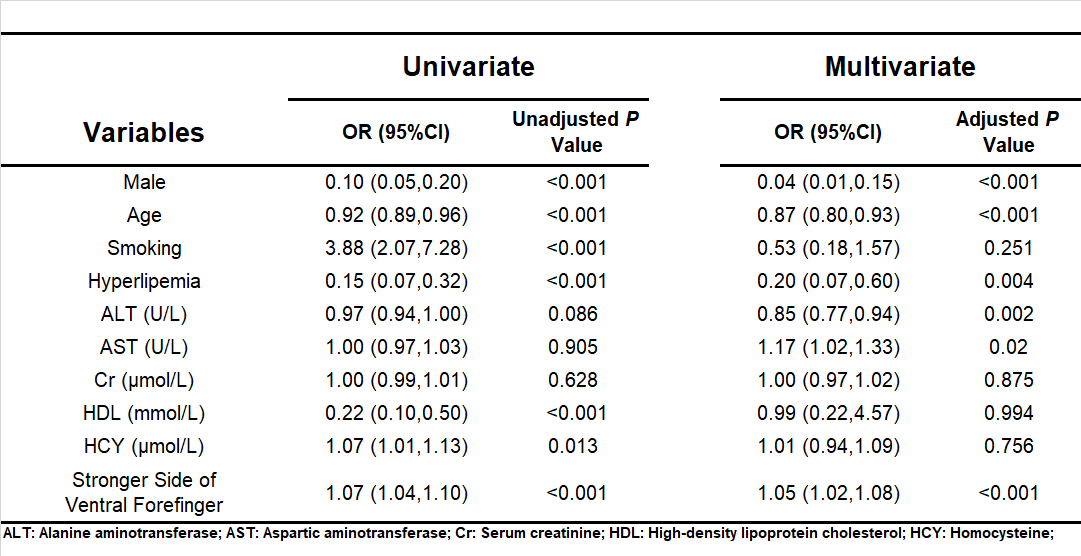

### Supplemental Table 4

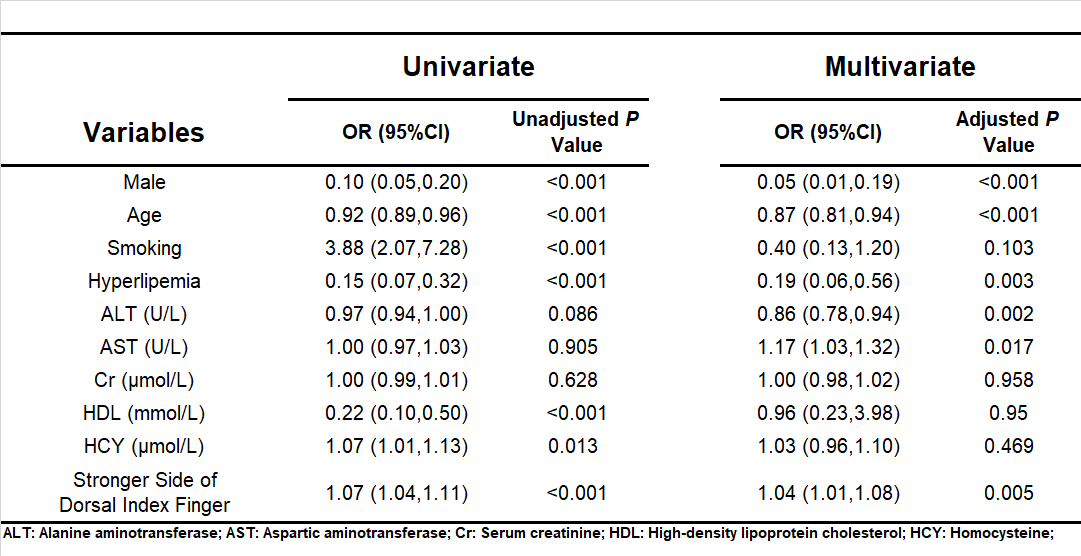

### Supplemental Table 5

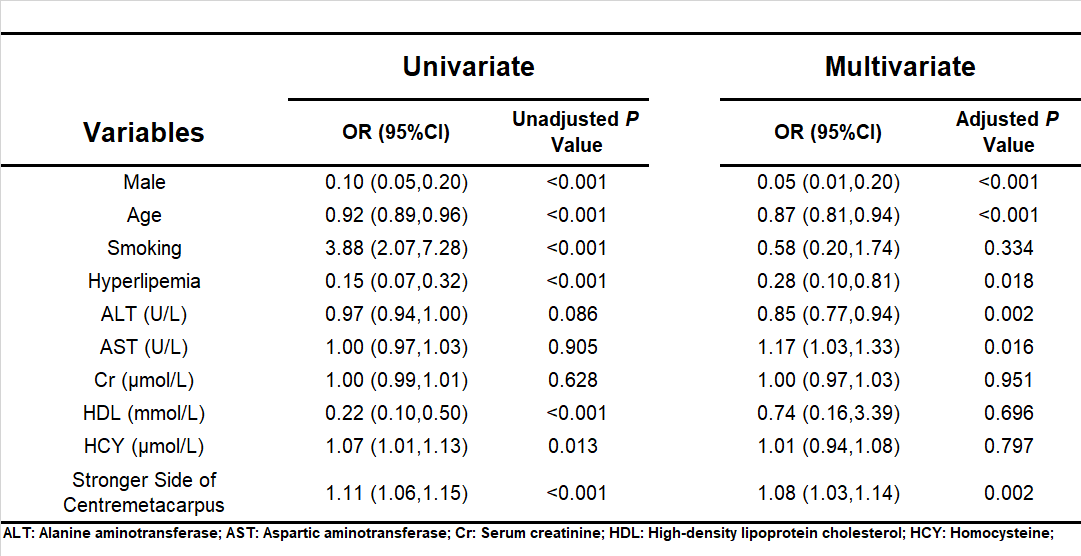

### Supplemental Table 6

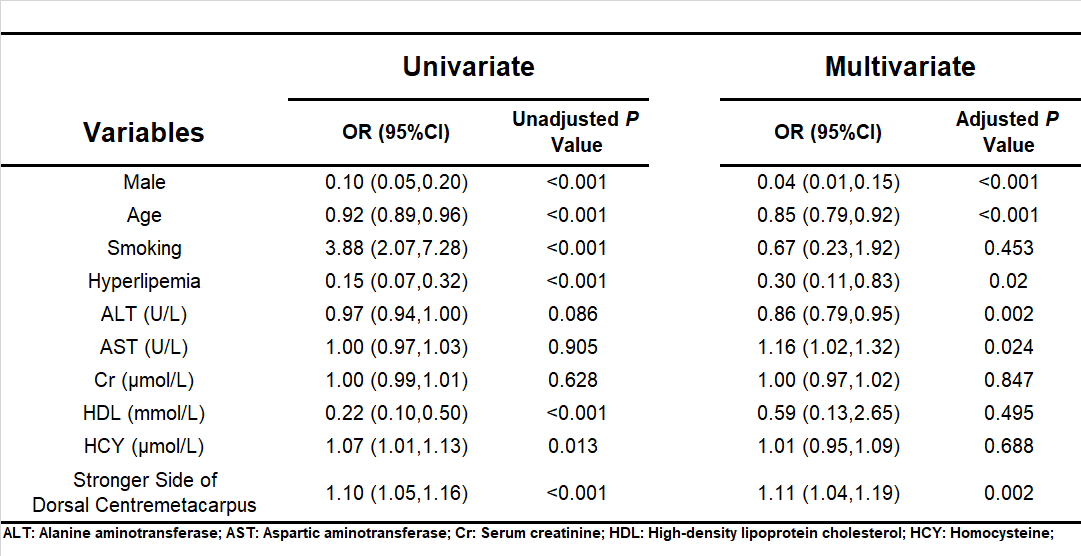

### Supplemental Table 7

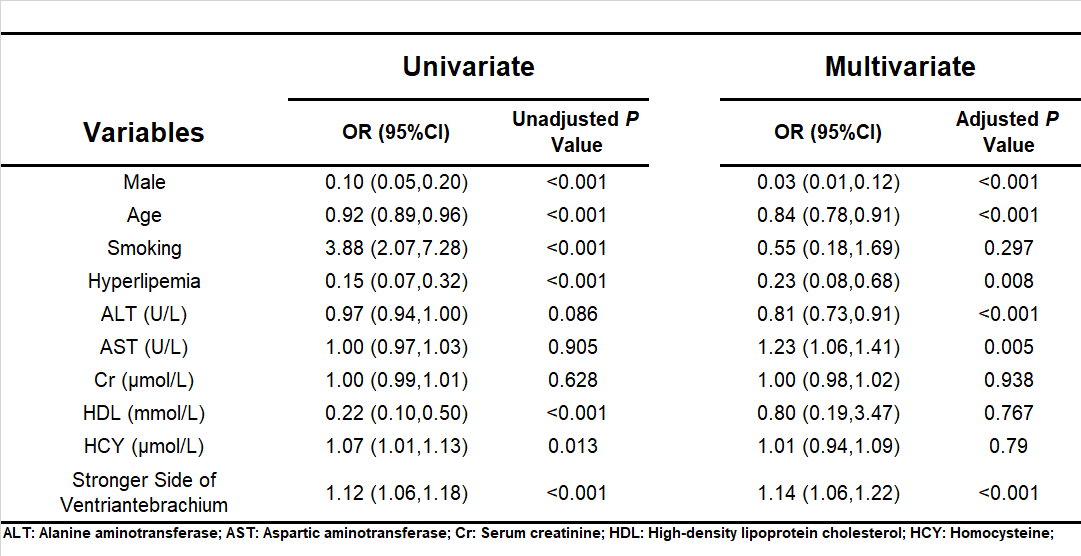

### Supplemental Table 8

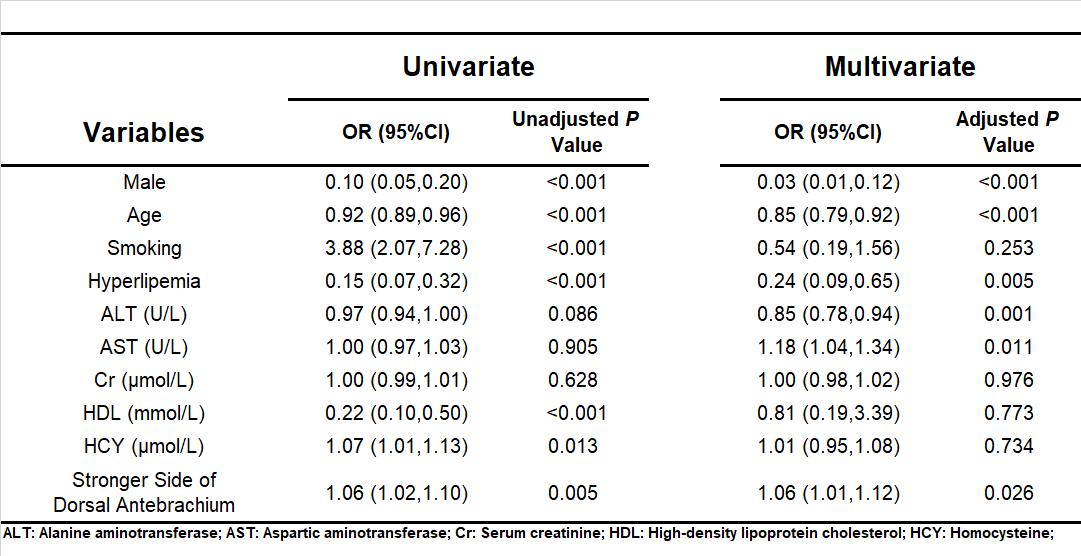

### Supplemental Table 9

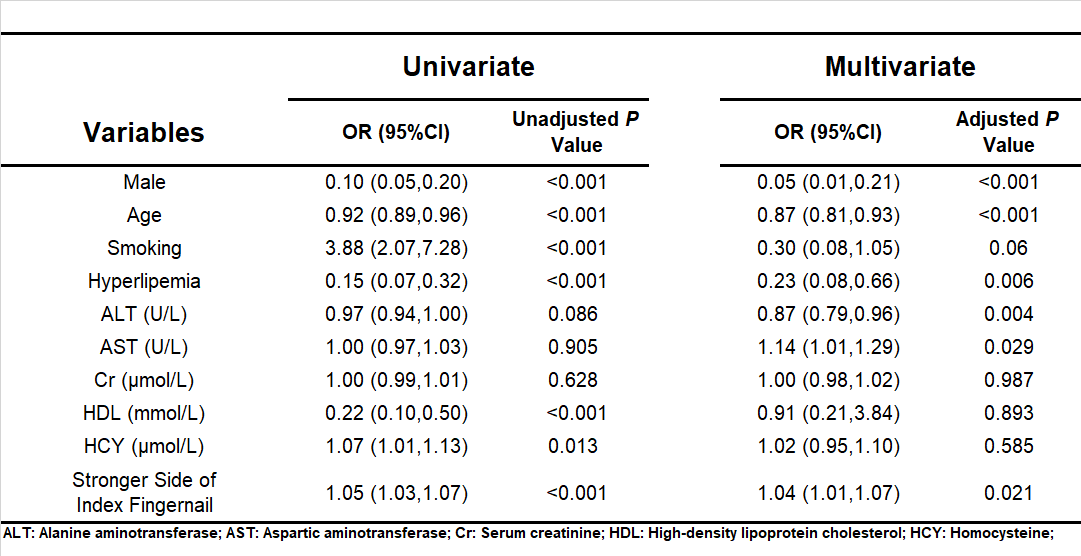

### Supplemental Table 10

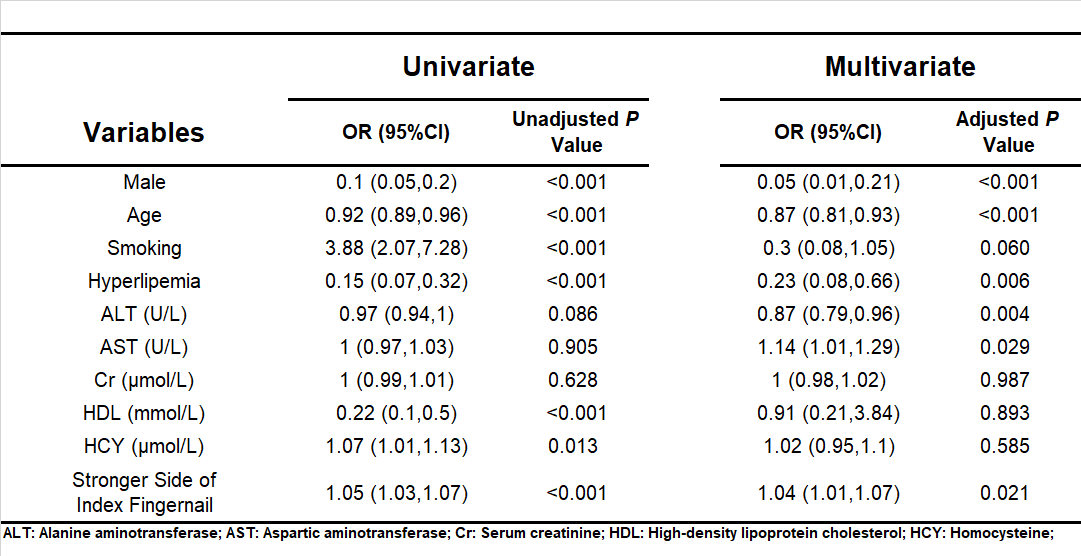

### Supplemental Table 12

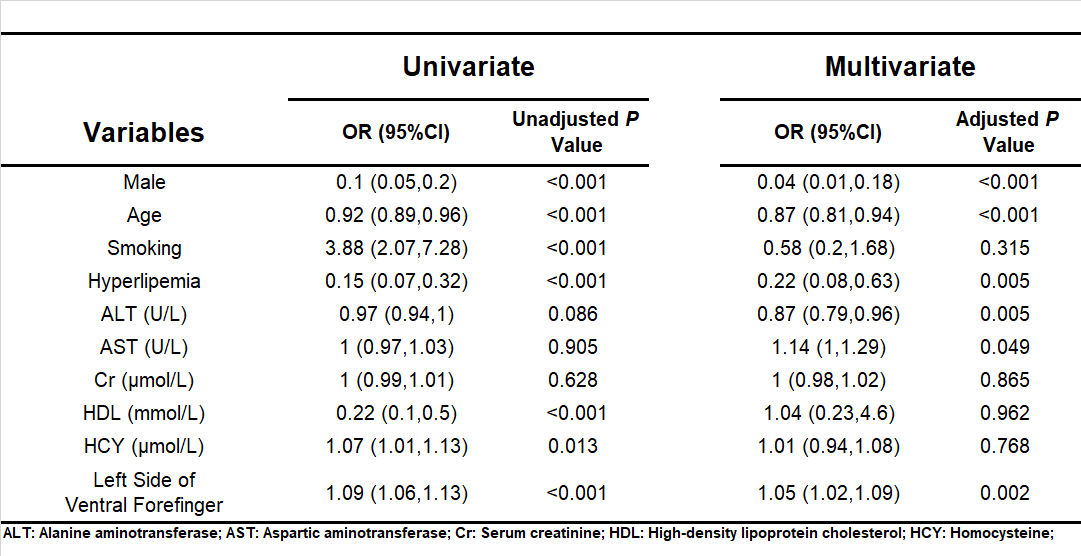

### Supplemental Table 13

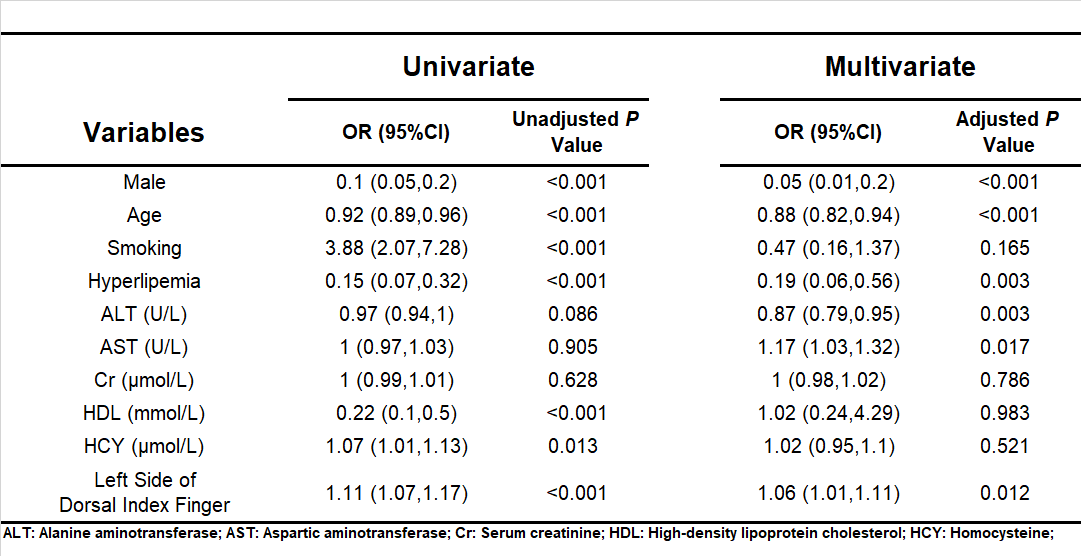

### Supplemental Table 14

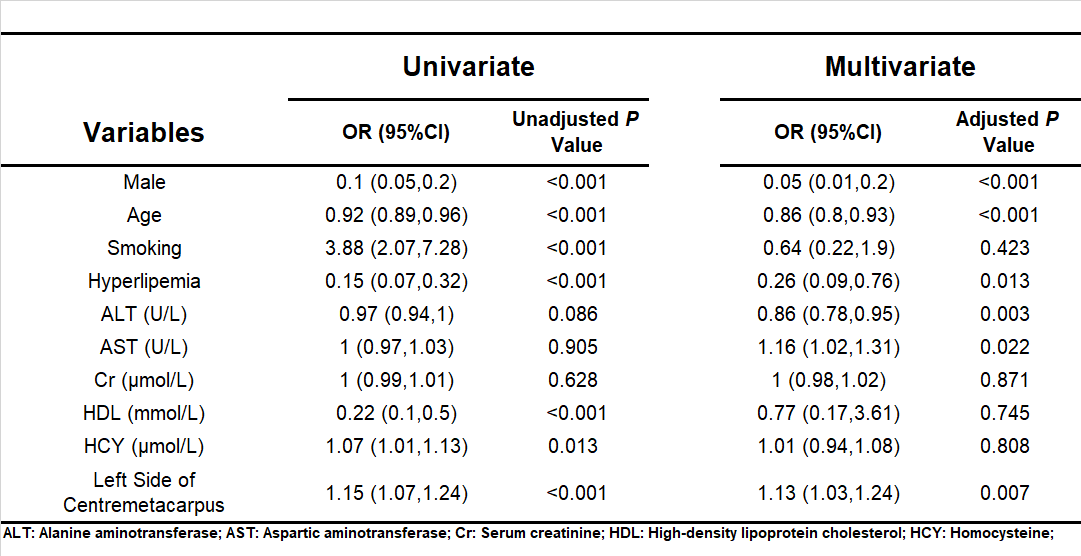

### Supplemental Table 15

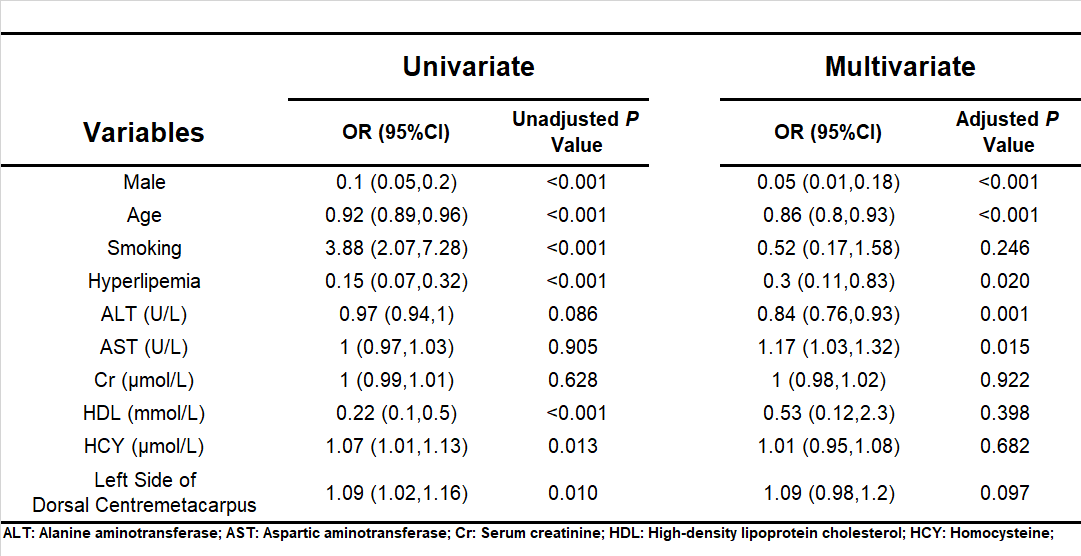

### Supplemental Table 16

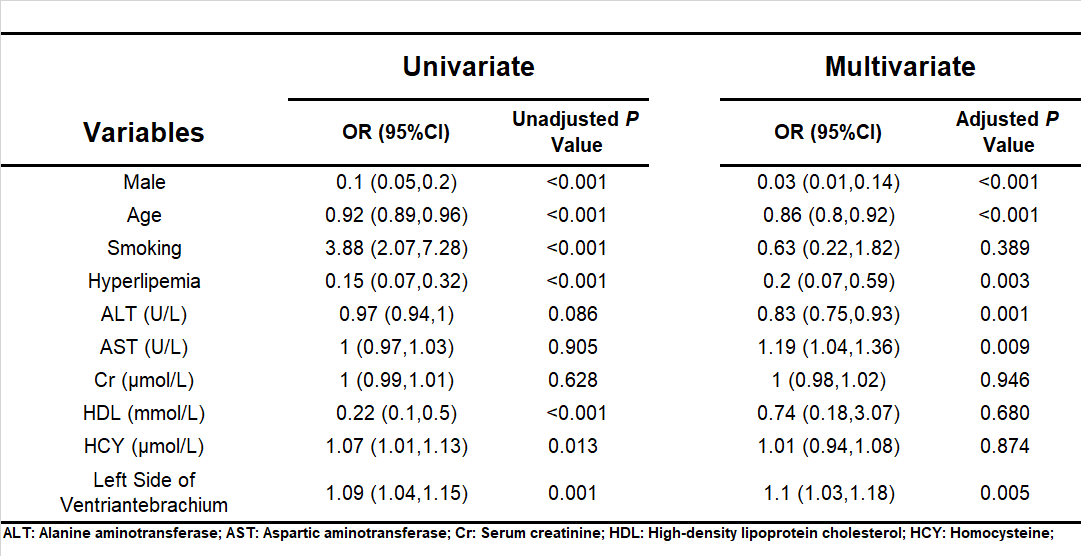

### Supplemental Table 17

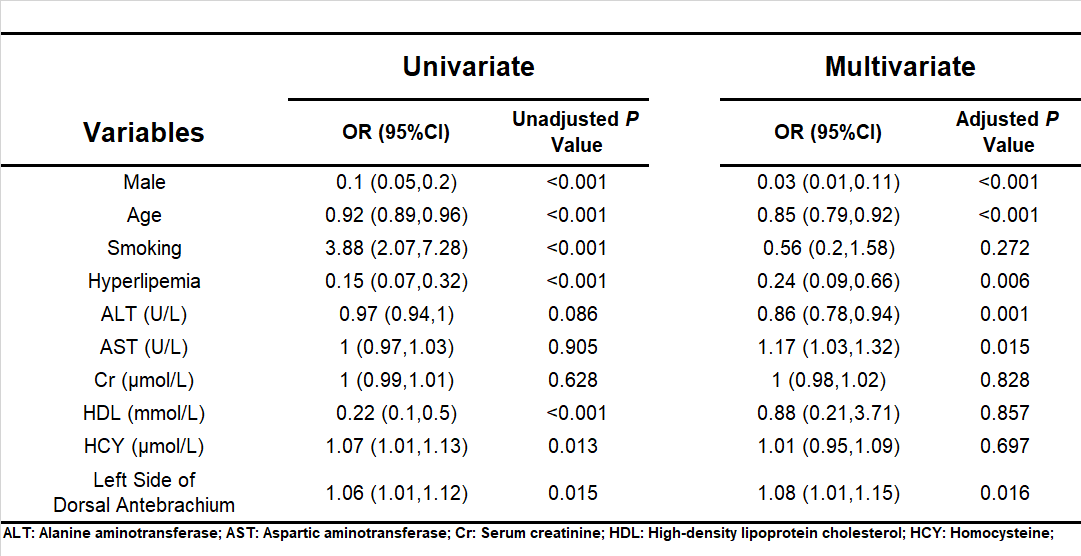

### Supplemental Table 18

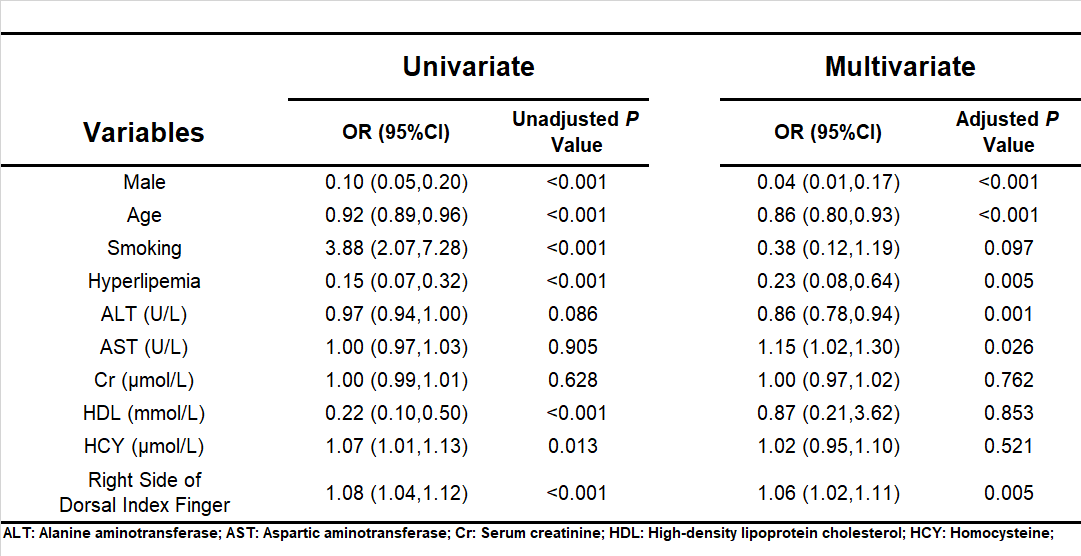

### Supplemental Table 19

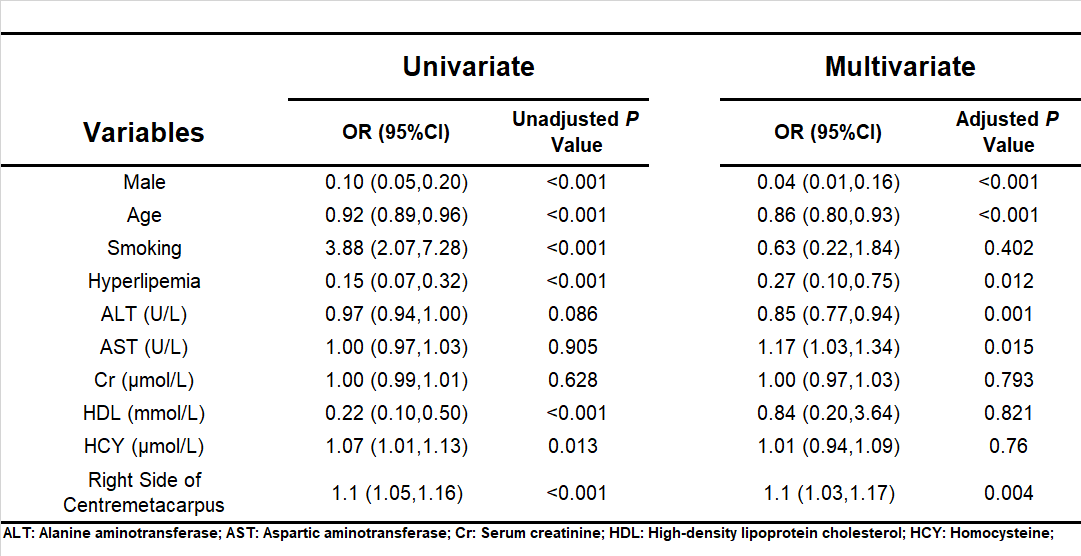

### Supplemental Table 20

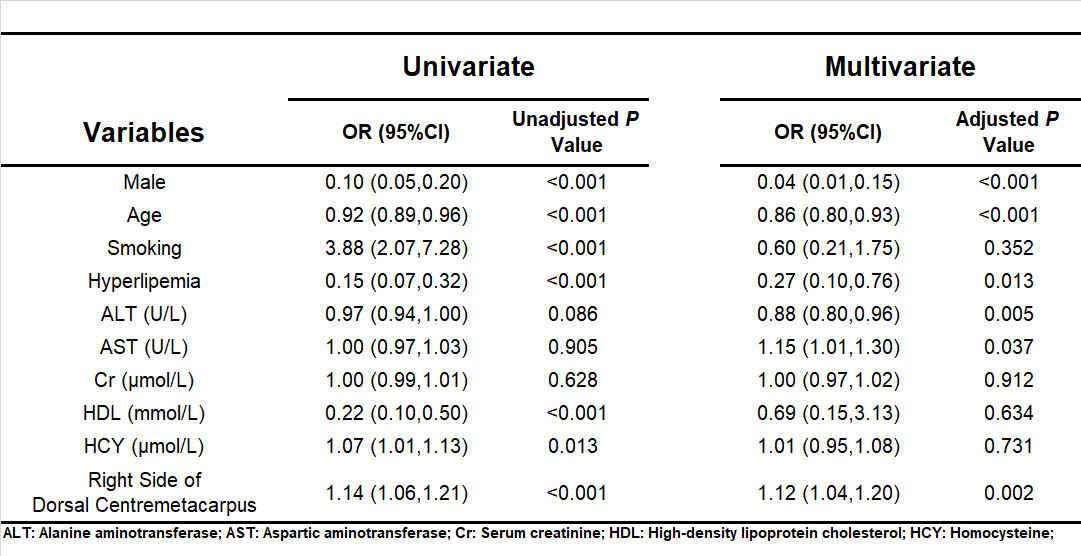

### Supplemental Table 21

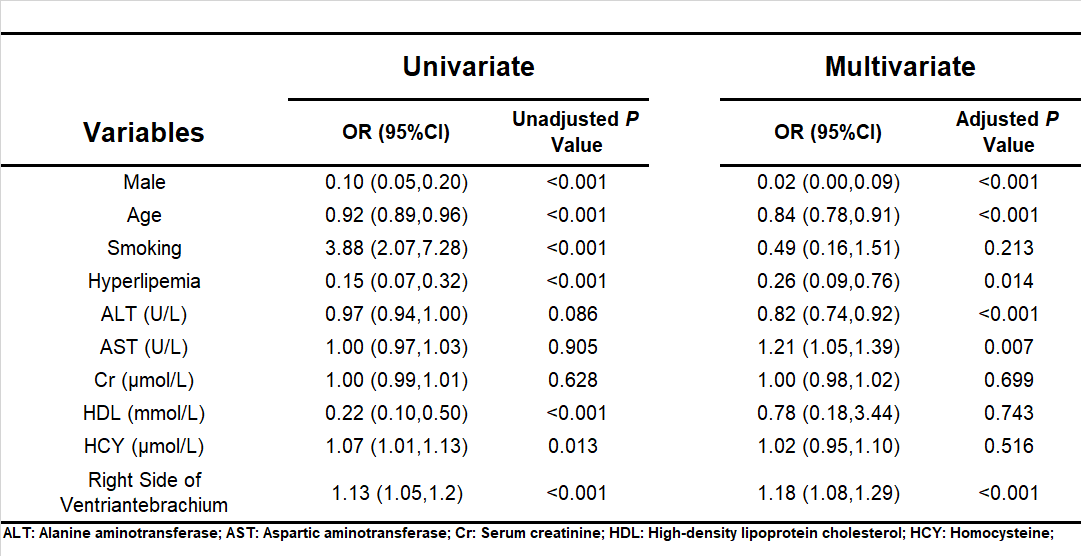

### Supplemental Table 22

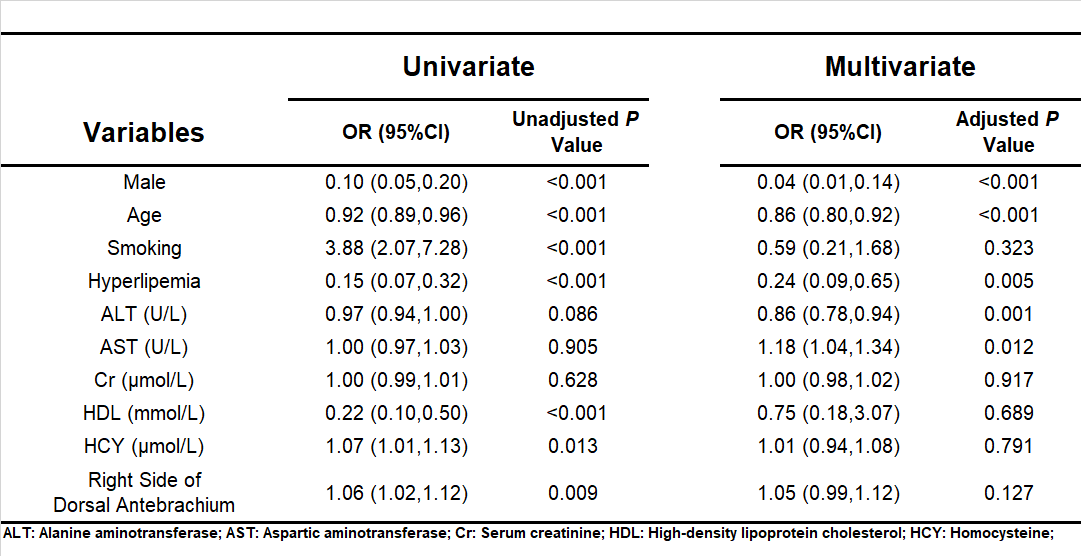

### Supplemental Table 23

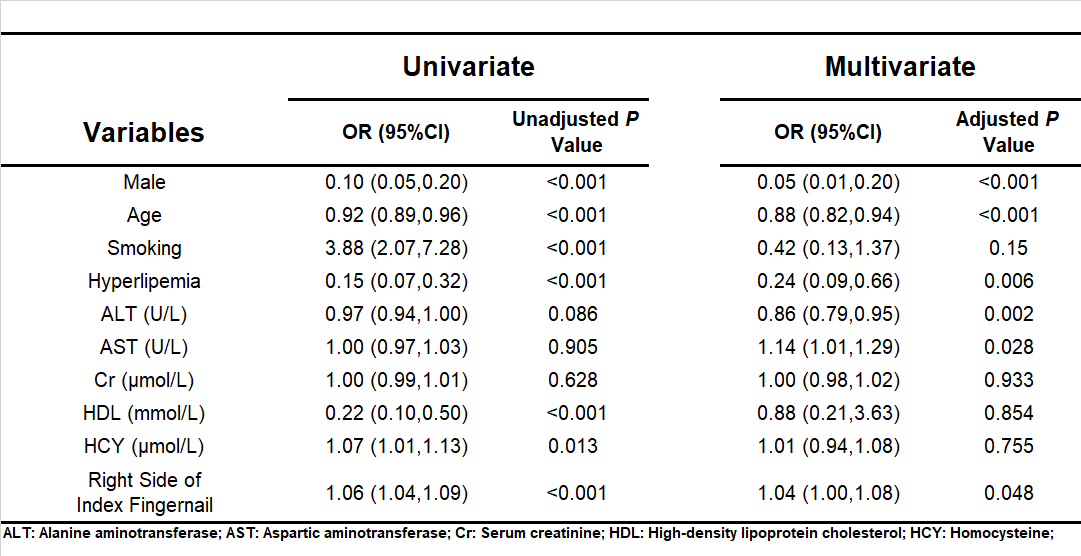

### Supplemental Table 24

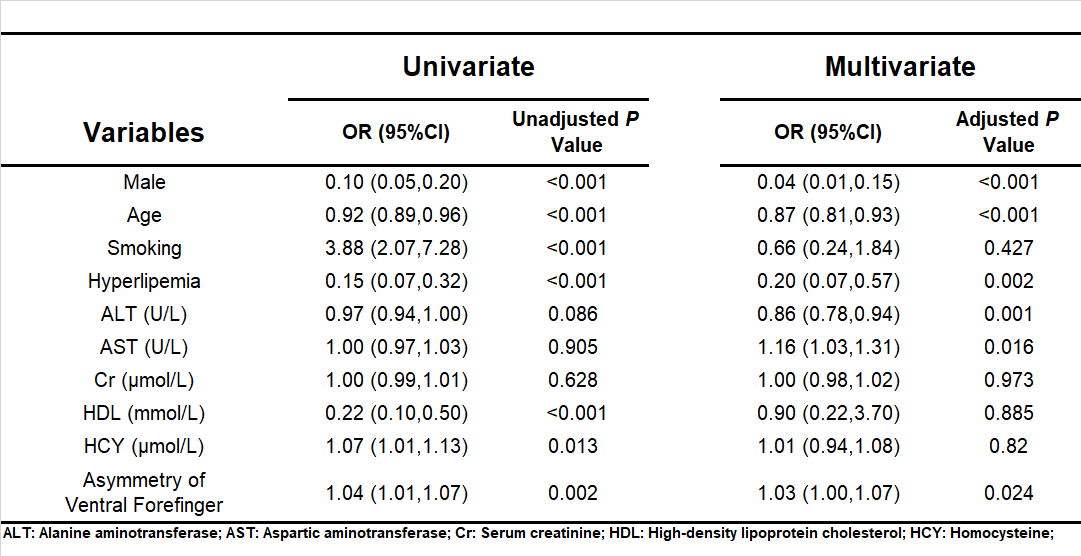

### Supplemental Table 25

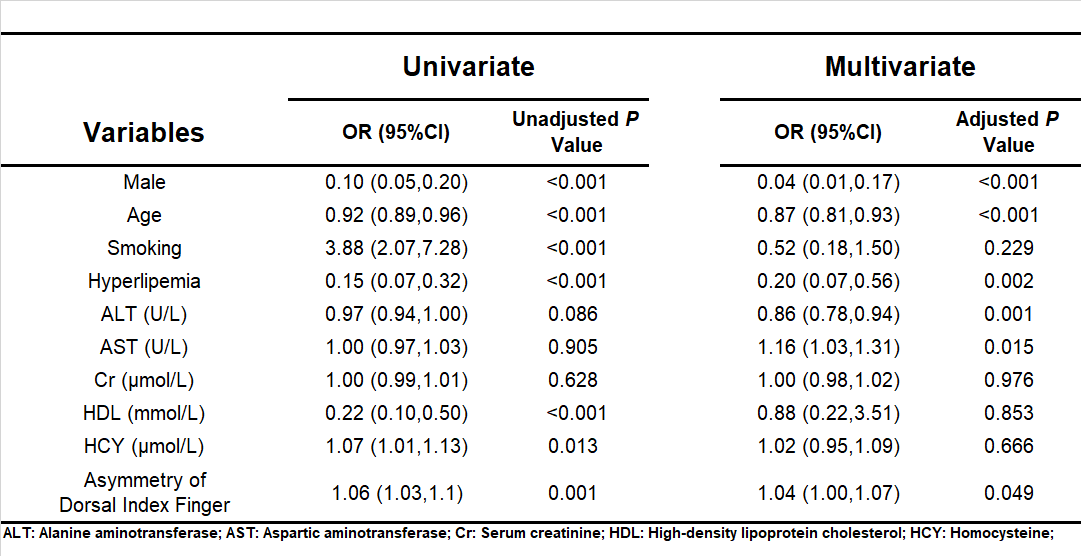

### Supplemental Table 26

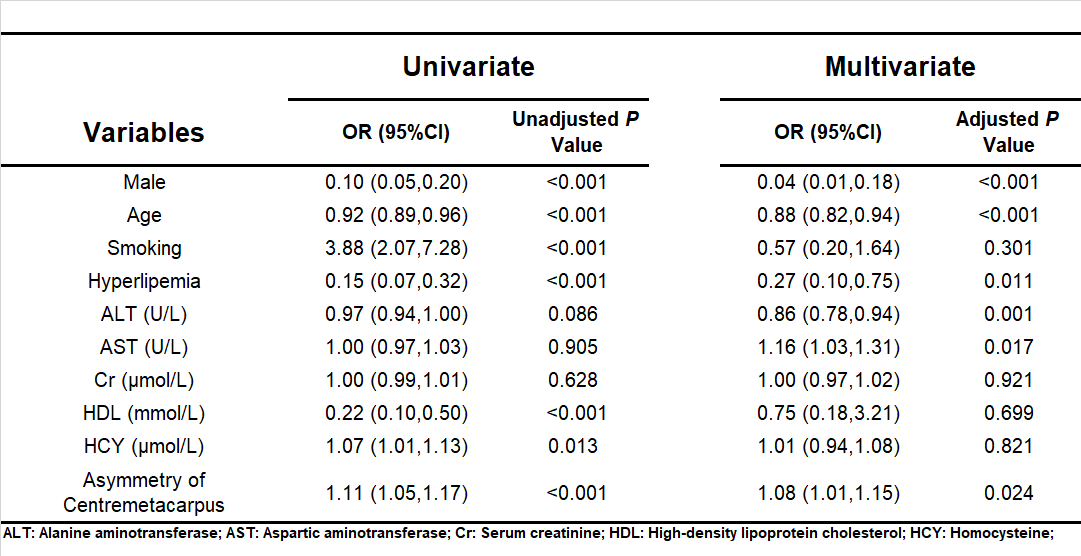

### Supplemental Table 27

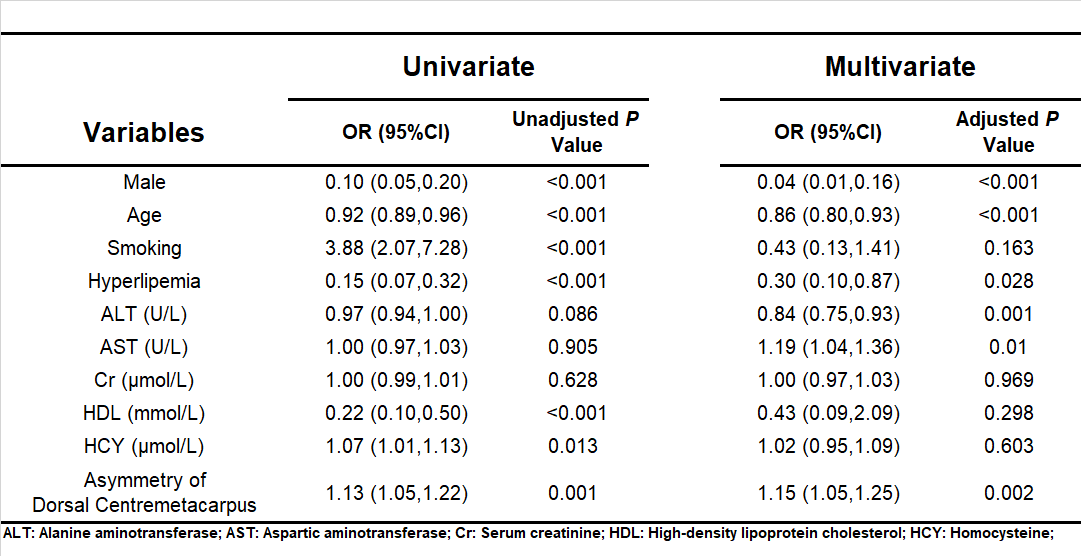

### Supplemental Table 28

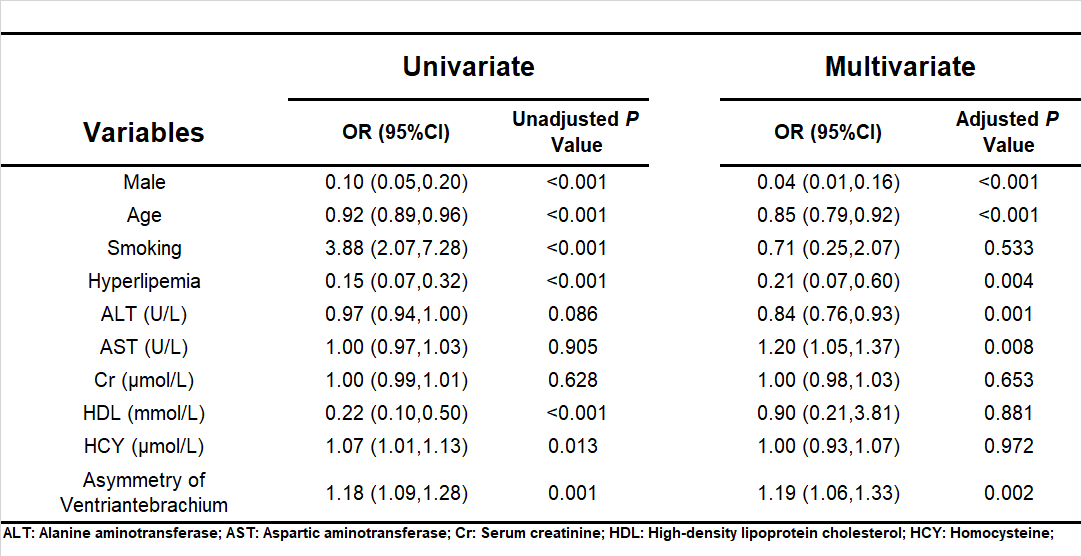

### Supplemental Table 29

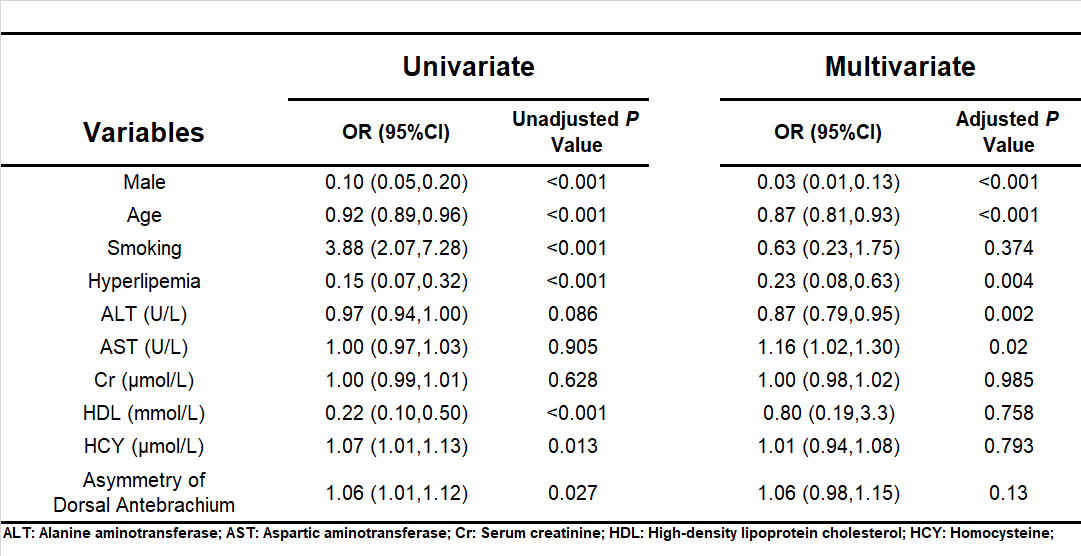

### Supplemental Table 30

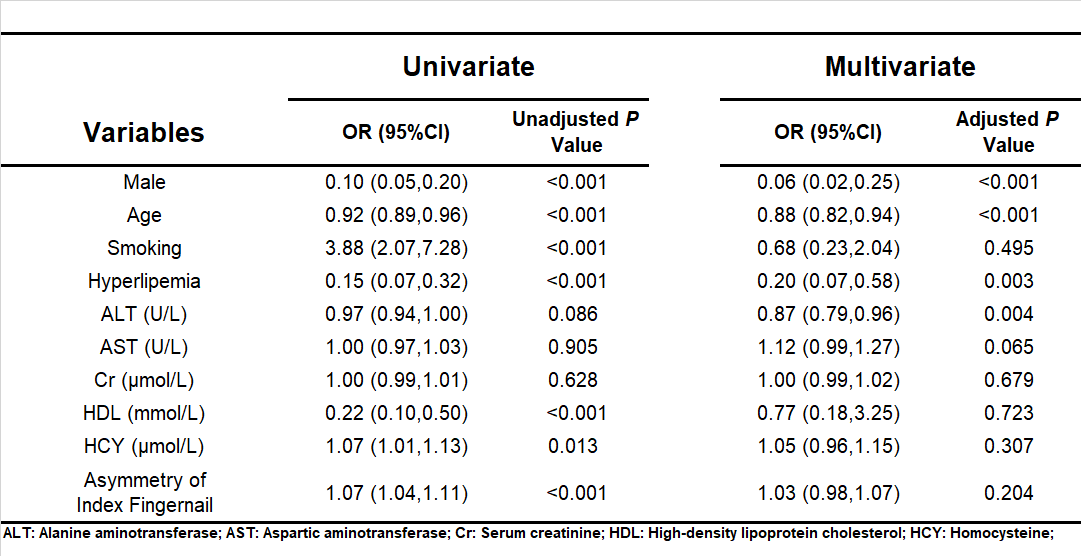

### Supplemental Table 31

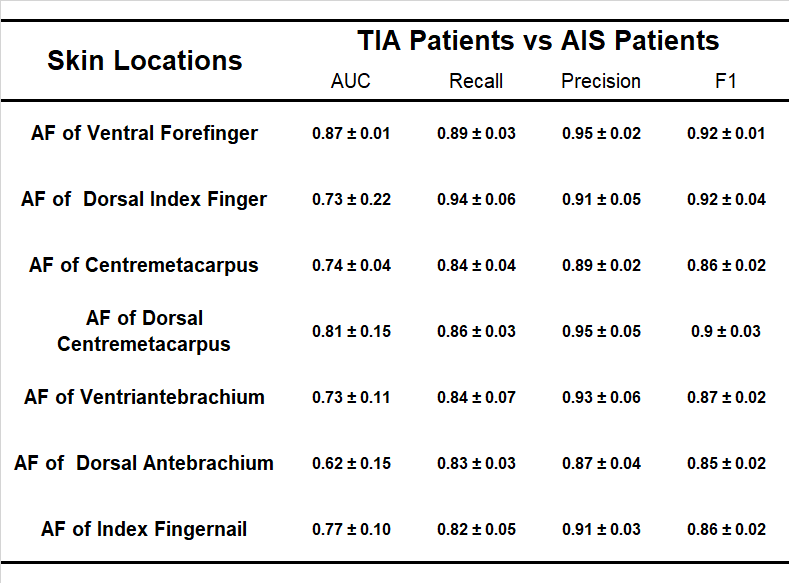

### Supplemental Table 32

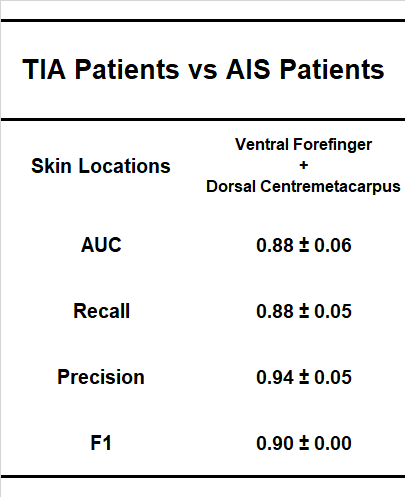
